# A fibroblast-rich transcriptomic axis reflects stromal remodeling within the CMS4 phenotype in colorectal cancer

**DOI:** 10.64898/2026.09.18.752659

**Authors:** Yuxi Zheng, Shuqiang Hao, Chonghe Jiang

## Abstract

Bulk-tumor expression signatures can reflect cellular composition as well as tumor-cell programs. We evaluated a frozen, equal-weight 87-gene score in colorectal cancer using 11 Gene Expression Omnibus (GEO) cohorts, TCGA COAD/READ, and E-MTAB-12862. In GEO (1,783 tumors; 441 progression-related events), the hazard ratio per score standard deviation was 1.33 (95% confidence interval 1.19–1.49). The estimate was similar in seven disease-free-survival cohorts. In 376 TCGA tumors, the hazard ratio was 1.25 (1.03–1.52) for progression-free interval. The association did not reproduce in E-MTAB-12862: among 948 stage I–III tumors with 436 recurrence-free-survival events, the hazard ratio was 1.05 (0.95–1.16). The frozen score correlated with CMS4, cancer-associated fibroblast, and stromal measures. The stromal-remodeling module produced a GEO estimate similar to the full score, whereas removing that module attenuated the association. Composition residualization did not detect additional prognostic information after accounting for the measured CMS4 and stromal components. Single-cell and spatial analyses localized the gene program to fibroblast-rich compartments. The 87-gene score mainly reflects a fibroblast-rich, matrix-remodeling component of CMS4 rather than an independent prognostic subtype. Its prognostic association varied across cohorts, and current evidence does not support clinical use.

## Introduction

Colorectal cancer (CRC) accounted for an estimated 1.9 million diagnoses and more than 900,000 deaths worldwide in 2022 [1]. Surgery remains the main curative treatment for localized colon cancer, with adjuvant chemotherapy selected according to stage and pathological risk. Treatment of metastatic disease combines systemic chemotherapy with molecularly selected targeted agents or immunotherapy and local treatment when appropriate [2, 3]. Clinical outcomes still vary among patients with tumors of similar anatomical stage.

Transcriptome-based classifications describe part of this heterogeneity. The consensus molecular subtype framework distinguishes immune, canonical, metabolic, and mesenchymal CRC states [5]. Bulk-tumor expression, however, combines signals from malignant, immune, vascular, and stromal cells. Variation in stromal content can affect both molecular classification and its apparent association with prognosis [6].

Cancer-associated fibroblasts (CAFs), extracellular matrix (ECM), endothelial cells, and myeloid cells influence invasion, immune exclusion, treatment response, and tissue remodeling [7]. Mesenchymal programs connect matrix remodeling with aggressive behavior. Regulators such as *ZEB1* and *ZEB2*, structural genes such as *VIM*, and several collagens participate in epithelial–mesenchymal plasticity or stromal biology [8]. An expression score enriched for these genes may therefore measure a malignant-cell state, microenvironmental composition, or a combination of the two.

Single-cell atlases can locate a bulk-derived program within specific cellular compartments. GSE178341 profiled 370,115 cells from CRC and adjacent tissue, and GSE132465 contains 63,689 cells from 23 patients, including ten matched normal-mucosa samples [9, 10]. Patient-level aggregation is needed when these data are used to compare tumor and normal tissue because the cells from one patient are not independent biological replicates.

Spatial transcriptomics can show whether a score varies across tissue regions, although Visium spots contain multiple cells and do not provide cell-type-specific measurements. GSE267401 contains matched primary CRC and liver-metastasis sections from two patients [11]. Ligand– receptor resources can support a focused expression screen in these datasets, but expression of both partners does not establish intercellular communication or causality [12].

We evaluated a neural–stromal remodeling program assembled from bladder-cancer and broader cancer literature before the present CRC outcome analyses. The primary analysis used a frozen 87-gene bulk score. A related 84-gene panel was used only for secondary biological annotation. We estimated the score’s association with progression-related outcomes in 11 GEO cohorts, transported the locked score to TCGA COAD/READ, and performed an independent external assessment in E-MTAB-12862 [4]. We separately examined overlap with CMS4 and stromal measures, module contributions, cellular localization, spatial distribution, and candidate ligand–receptor expression (Fig. 1) to determine what biological state the score measured. The score was not developed through a CRC-specific systematic review, and the original GEO split was selected during an exploratory search. We therefore treat it as a retrospective research score rather than a clinical biomarker.

**Figure 1.**
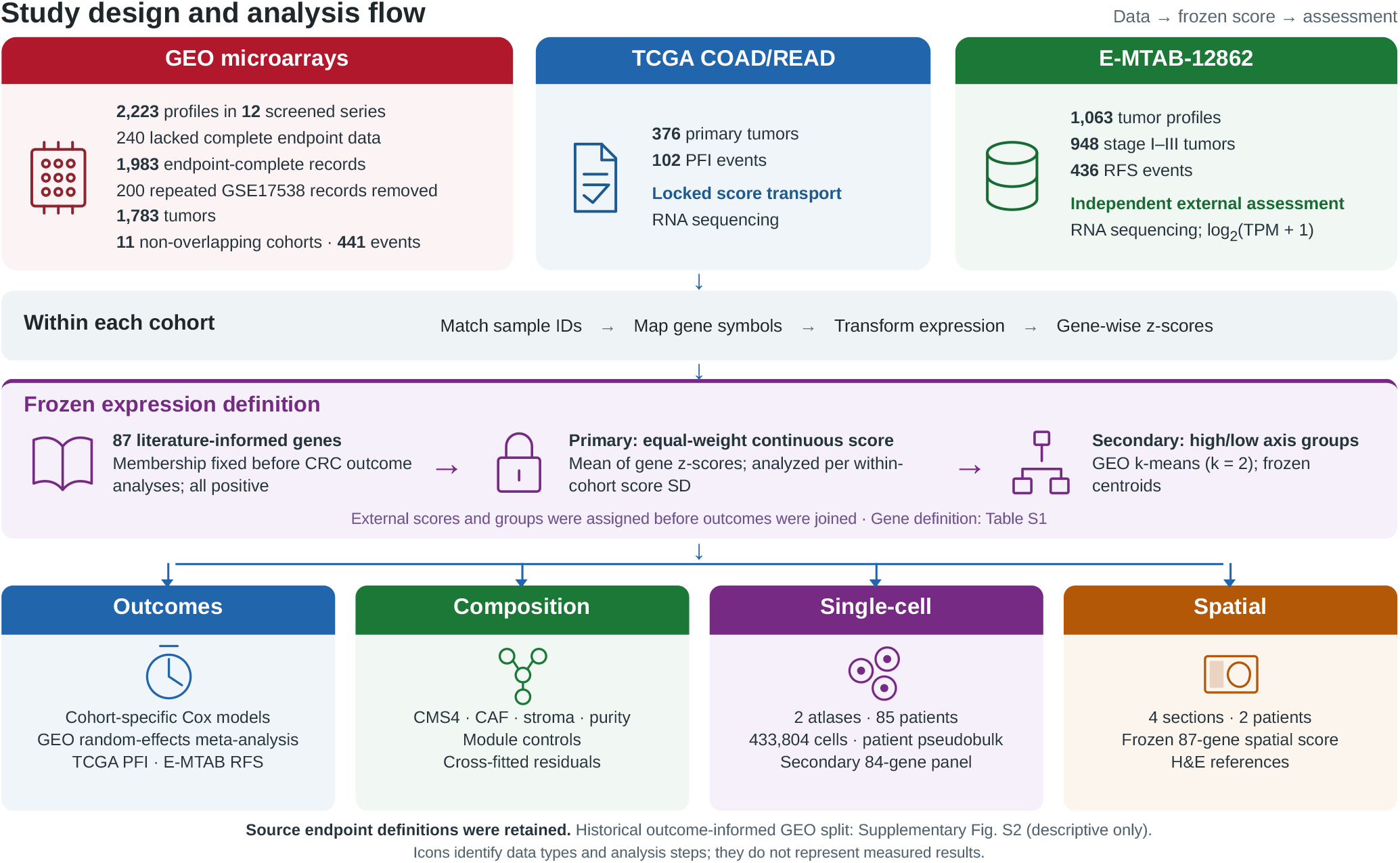
Study design and analysis flow. Public expression profiles were matched to clinical follow-up, checked for duplicate records, mapped to gene symbols, and standardized within each cohort. The literature-informed 87-gene list was fixed before the CRC outcome analyses. Its equal-weight continuous score was the primary exposure; locked high- and low-axis groups were secondary. Cohort-specific models retained the original DFS, RFS, MFS, MRFS, PFI, or PFS definition and addressed the shared progression-related event-free survival question. GEO estimates were pooled by random-effects meta-analysis, and the score was transported without outcome-based modification to TCGA and E-MTAB-12862. Module, composition, single-cell, and spatial analyses examined the biological source of the score. The historical outcome-informed GEO partition is shown only in Supplementary Fig. S2. CAF, cancer-associated fibroblast; PFI, progression-free interval; RFS, recurrence-free survival.

## Materials and Methods

### Study datasets and design

We performed a secondary analysis of public CRC transcriptomic and clinical datasets (Table 1). We screened 12 GEO series generated primarily on the Affymetrix Human Genome U133 Plus 2.0 Array (GPL570) [14]. Eleven non-overlapping cohorts contributed 1,783 expression-matched tumors and 441 progression-related events to the primary meta-analysis. The 200 endpoint-complete GPL570 records in GSE17538 duplicated records already present in GSE17536 and GSE17537 and were excluded before modeling. TCGA COAD/READ RNA-sequencing and clinical data were obtained through UCSC Xena; 376 primary tumors with 102 progression-free-interval (PFI) events were eligible for locked transport assessment. E-MTAB-12862 provided a separate population-based series of 1,063 primary CRC tumors with gene-symbol transcript-per-million (TPM) expression and long-term follow-up [4]. Its primary recurrence-free-survival (RFS) analysis included 948 stage I–III tumors and 436 events.

**Table 1.**
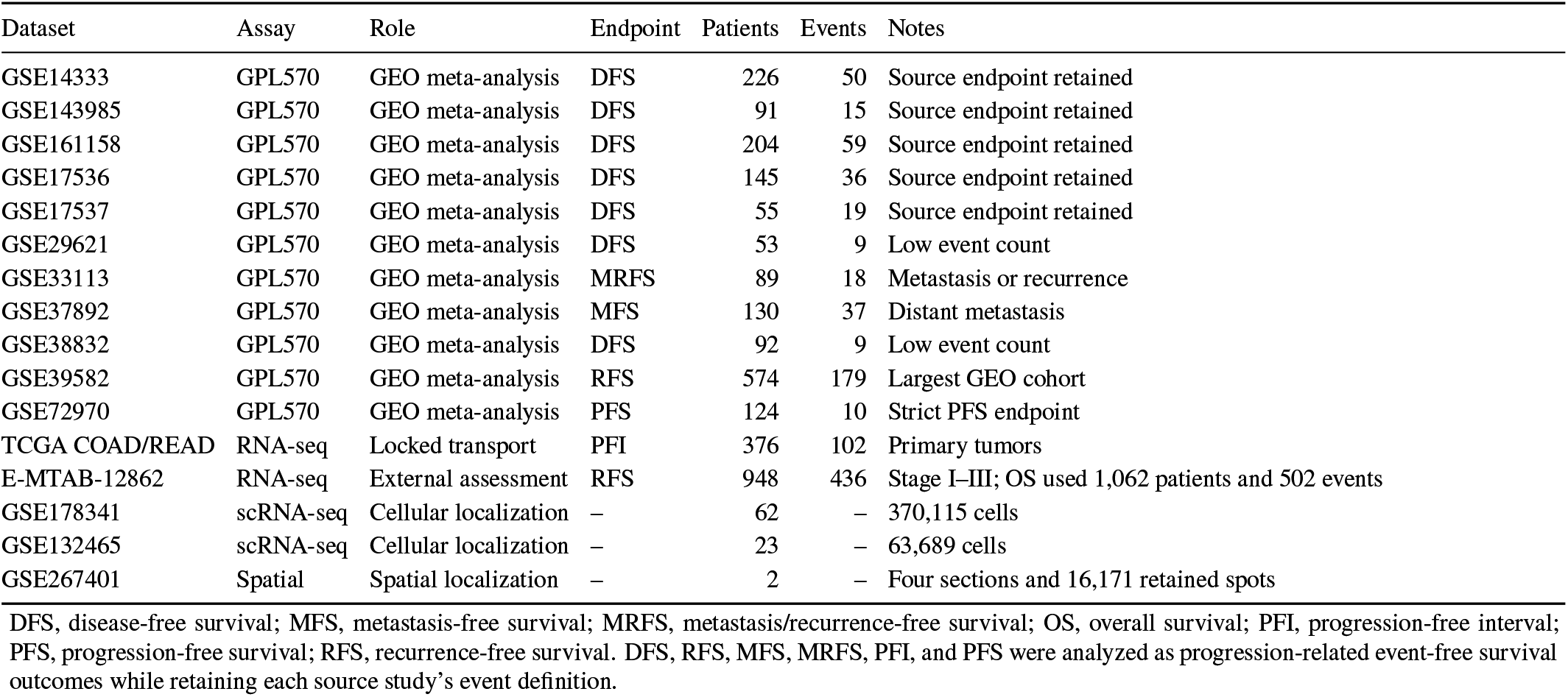
Datasets included in the study. Patient and event counts refer to the analysis listed in the Role column.

| Dataset | Assay | Role | Endpoint | Patients | Events | Notes |
| --- | --- | --- | --- | --- | --- | --- |
| GSE14333 | GPL570 | GEO meta-analysis | DFS | 226 | 50 | Source endpoint retained |
| GSE143985 | GPL570 | GEO meta-analysis | DFS | 91 | 15 | Source endpoint retained |
| GSE161158 | GPL570 | GEO meta-analysis | DFS | 204 | 59 | Source endpoint retained |
| GSE17536 | GPL570 | GEO meta-analysis | DFS | 145 | 36 | Source endpoint retained |
| GSE17537 | GPL570 | GEO meta-analysis | DFS | 55 | 19 | Source endpoint retained |
| GSE29621 | GPL570 | GEO meta-analysis | DFS | 53 | 9 | Low event count |
| GSE33113 | GPL570 | GEO meta-analysis | MRFS | 89 | 18 | Metastasis or recurrence |
| GSE37892 | GPL570 | GEO meta-analysis | MFS | 130 | 37 | Distant metastasis |
| GSE38832 | GPL570 | GEO meta-analysis | DFS | 92 | 9 | Low event count |
| GSE39582 | GPL570 | GEO meta-analysis | RFS | 574 | 179 | Largest GEO cohort |
| GSE72970 | GPL570 | GEO meta-analysis | PFS | 124 | 10 | Strict PFS endpoint |
| TCGA COAD/READ | RNA-seq | Locked transport | PFI | 376 | 102 | Primary tumors |
| E-MTAB-12862 | RNA-seq | External assessment | RFS | 948 | 436 | Stage I–III; OS used 1,062 patients and 502 events |
| GSE178341 | scRNA-seq | Cellular localization | – | 62 | – | 370,115 cells |
| GSE132465 | scRNA-seq | Cellular localization | – | 23 | – | 63,689 cells |
| GSE267401 | Spatial | Spatial localization | – | 2 | – | Four sections and 16,171 retained spots |

DFS, disease-free survival; MFS, metastasis-free survival; MRFS, metastasis/recurrence-free survival; OS, overall survival; PFI, progression-free interval; PFS, progression-free survival; RFS, recurrence-free survival. DFS, RFS, MFS, MRFS, PFI, and PFS were analyzed as progression-related event-free survival outcomes while retaining each source study’s event definition.

Cellular localization used GSE178341, which contained 370,115 retained cells from 62 patients, and GSE132465, which contained 63,689 cells from 23 patients [9, 10]. The spatial analysis used four GSE267401 sections from two patients, with one primary CRC and one liver-metastasis section per patient. After quality control, 16,171 spatial spots remained.

### GEO cohort endpoint harmonization

Processed expression matrices, platform annotations, and clinical metadata were standardized separately within each GEO dataset. GPL570 probes were mapped to gene symbols, and probes mapping to the same symbol were averaged. The source studies reported DFS, RFS, MFS, metastasis/recurrence-free survival, or PFS. Follow-up times were converted to months, and each cohort retained its source endpoint definition and event coding.

These endpoints address a shared clinical question: how long a patient remains free of a source-defined disease event after the study baseline. We therefore refer to them collectively as *progression-related event-free survival*, or *PFS-like survival* in descriptions of the historical workflow. An event could be recurrence, metastasis, progression, a new tumor event, or death, depending on the original study. This harmonization provides a common analysis label and time unit; it does not redefine DFS, RFS, MFS, MRFS, or PFI as strict PFS. The exploratory clustering workflow joined the standardized time and event fields for its original Kaplan–Meier displays. The primary analysis did not pool individual patient records under one endpoint. Cox models were fitted separately in each cohort, and the cohort log hazard ratios were combined by meta-analysis. We also analyzed the seven DFS cohorts alone and repeated the meta-analysis after removing each singleton non-DFS endpoint family.

### Construction of the frozen 87-gene transcriptomic axis

The primary axis was a predefined 87-gene set assembled before the CRC survival analyses from literature on stromal remodeling, mesenchymal biology, neural guidance, and related tumor programs. Gene membership and direction were not changed according to CRC outcomes. Supplementary Table S1 lists all 87 genes and their assigned modules.

Within each cohort, every gene was standardized using the cohort mean and population standard deviation. The continuous score was the unweighted mean of the 87 gene-wise z-scores, with all genes contributing in the positive direction. PCA and *k*-means were not part of this score definition. The continuous score was then standardized within cohort before Cox regression, so each hazard ratio represents a one-standard-deviation increase in that cohort’s score.

For module analyses, the 87 genes were assigned to five non-overlapping broad categories: stromal remodeling (*n* = 55), neural-guidance signaling (*n* = 15), neural-related phenotype control (*n* = 8), neural-related signaling (*n* = 5), and neural-growth-factor-like signaling (*n* = 4). These categories describe gene membership; they are unrelated to the leave-one-cohort-out analyses.

A related 84-gene panel was retained for secondary annotation and single-cell analyses. It shares 83 genes with the primary score, excludes *ERBB2, ERBB3, ERBB4*, and *NRG1*, and adds *EGFR*. The four excluded genes were measurable in the bulk datasets, so the difference does not reflect expression availability. The available provenance records do not document why this membership change was made. The panel includes predefined directions for direction-aware analyses. We used its membership without directions to calculate the descriptive *84-gene localization score* and used its directions to calculate the patient-level *84-gene signed state score*. Neither secondary score replaces or modifies the frozen 87-gene primary score.

### Exploratory unsupervised clustering

PCA was applied to the evidence-gene matrix after expression values were standardized within each GEO dataset. Survival outcomes were not used in PCA. K-means clustering was then applied to the PCA representation. The original analysis used *k* = 2, with *k* = 3 examined as a sensitivity analysis. The cluster with higher aggregate expression of the axis genes was labeled high axis.

The source workflow evaluated all six-versus-six dataset partitions and selected split 595 using both expression separation and PFS-like survival behavior. Survival was not an input to PCA or *k*-means, but it influenced which dataset partition was reported. This outcome-informed selection creates selection bias in the subsequent Kaplan–Meier comparison. The labels *discovery* and *internal assessment* describe the two arms of the selected partition; neither arm is an independent validation set, and their log-rank *p* values are retained only to document the exploratory analysis [15].

### Primary survival analysis and cohort-level meta-analysis

The frozen continuous score was evaluated with a separate Cox proportional-hazards model in each cohort [16]. Cohort log hazard ratios were combined by random-effects meta-analysis using restricted maximum likelihood to estimate τ^2^ and Knapp–Hartung confidence intervals. Leave-one-cohort-out analyses assessed the influence of each study. The endpoint family was fixed in the cohort manifest as DFS, RFS, MFS, metastasis/recurrence-free survival, or PFS. We repeated the analysis in the seven DFS cohorts and after excluding each singleton RFS, MFS, or metastasis/recurrence-free cohort. We did not fit endpoint meta-regression because four of the five endpoint categories contained only one cohort.

A separate leave-one-dataset-out classifier analysis retrained the expression scaler and *k*-means classifier without the held-out cohort. Survival data from that cohort were joined only after expression-based group assignment. Cohorts with few events remained in the meta-analysis, and their imprecision is reflected in their confidence intervals.

### Differential expression and enrichment

High-versus low-axis differential expression used empirical-Bayes linear modeling in `limma` [20]. The genome-wide analysis tested 22,834 genes. Benjamini–Hochberg FDR controlled multiple testing [22]. Results were summarized at FDR below 0.05, with absolute log-fold-change thresholds of 0.5 for the principal display and 1.0 for a more stringent count. A selected-gene analysis separately assessed members of the secondary 84-gene panel.

Ranked differential-expression results were evaluated against Hallmark, Gene Ontology Biological Process, KEGG, and custom axis modules using gene-set enrichment analysis and hypergeometric over-representation tests [23, 24]. FDR was applied within each enrichment family.

### Immune and stromal inference

EPIC estimated cancer, immune, endothelial, and CAF fractions from normalized bulk expression [25]. Additional marker-based standardized scores represented fibroblast, macrophage, endothelial, stromal, and immune programs; these were exploratory and were not treated as validated deconvolution estimates. High- and low-axis distributions were compared with Wilcoxon rank-sum tests, and rank-biserial correlations summarized effect direction and magnitude.

### Locked score transport to TCGA COAD/READ

TCGA COAD/READ PanCanAtlas expression, phenotype, and survival data were obtained through UCSC Xena [26]. Primary tumors were matched across files. PFI definitions followed the TCGA Clinical Data Resource, with event 1 indicating progression, recurrence, a new tumor event, or death and event 0 indicating censoring [27]. Time was converted from days to months. Each of the 87 genes was standardized across TCGA primary tumors, and their unweighted mean was standardized again to define the continuous score. The frozen GEO scaler, *k*-means centroids, and cluster orientation were used only to assign the locked high- and low-axis groups. Both scores were calculated before TCGA survival outcomes were joined. Continuous-score and locked high-versus-low Cox models were then fitted in the 376 PFI-evaluable tumors.

### Frozen-score external assessment and exploratory model development in E-MTAB-12862

E-MTAB-12862 was analyzed under a locked protocol using the official gene-symbol TPM matrix and public clinical supplement [4]. Scale inspection was completed without survival outcomes and showed raw TPM-scale values, so expression was transformed as log_2_ (TPM + 1). Duplicate symbols were averaged, the frozen *CXCR7*-to-*ACKR3* alias was applied, and all 87 genes were available. Each gene was standardized across the 1,063 tumor samples, after which the equal-weight score and locked group assignment were calculated without reference to outcome.

The primary endpoint was RFS among 948 stage I–III patients, with 436 events. The source study defined RFS from surgery to local or distant recurrence or death, with censoring at last follow-up. Overall survival (OS) among 1,062 patients, with 502 deaths, was secondary. The primary model was RFS ~ axis, with the axis expressed per E-MTAB population standard deviation. Secondary analyses included categorical stage adjustment, the locked group comparison, proportional-hazards diagnostics, module scores, a *VIM* control, and correlations with the source CMS4 label and composition proxies.

After the locked analyses were completed, the same 948 stage I–III tumors were used for exploratory model development. A least absolute shrinkage and selection operator (LASSO)-penalized Cox model included the standardized expression values of all 87 genes. The penalty was selected by five-fold cross-validation in E-MTAB-12862, and the model was then fitted to the full cohort. Thirty-four genes had nonzero coefficients. The resulting linear predictor was standardized within the RFS analysis set.

We estimated the full-cohort association between this score and RFS and calculated the concordance index. For Kaplan– Meier visualization, the highest and lowest score quartiles were compared, with 237 patients in each group; the middle 50% of patients were excluded only from this comparison. Fold-specific concordance indices were also calculated after refitting the coefficients in each training fold. The penalty had already been selected using the full E-MTAB-12862 cohort, so this assessment was not nested cross-validation. The LASSO estimates are therefore reported as same-cohort model-development results and not as independent validation.

### CMS4/stroma comparator analyses

CMS4-like signal was measured with CMScaller distance-derived affinity, which was treated as a continuous affinity rather than a posterior probability. EPIC CAF fractions were used for GEO where available; the corresponding TCGA and unavailable-resource measures were explicitly labeled marker proxies. Within each cohort, Spearman correlations quantified overlap between the frozen score and CMS4 affinity, CAF/proxy, stroma proxy, and purity proxy. Nested Cox comparisons used stage, CMS4 affinity, and one stromal comparator at a time, with the axis added only in the expanded model. Collinearity and low-event warnings were retained rather than resolved by unplanned cohort removal.

For the cross-fitted residual analysis, the four composition predictors were standardized within cohort without using outcome data. Each GEO cohort was held out in turn; a linear nuisance model of the frozen score on CMS4 affinity, CAF, stroma, and purity was fitted in the other ten cohorts and applied to the held-out samples. Because the original axis was already expressed per within-cohort standard deviation, the observed-minus-predicted residual remained on that original-axis scale and was not re-standardized. It was then entered into a univariable Cox model. A prespecified ridge nuisance model, tuned only among training cohorts, was used as a sensitivity analysis. Held-out log-hazard ratios were pooled with REML and Knapp–Hartung uncertainty. For external assessment, the nuisance model was trained on all GEO cohorts and applied once to TCGA. Survival time and event status were never inputs to residual construction.

For the stage-adjusted sensitivity, cohorts were eligible only when their frozen analysis table contained usable stage coding. Within each of seven eligible cohorts, separate Cox models included categorical stage with either the original axis or the cross-fitted residual axis. No stage value was imputed, and no cohort was removed according to its result. Cohort log-hazard ratios were pooled with the same REML and Knapp–Hartung procedure.

### Simulation benchmark

The frozen simulation design preserved the 11 GEO cohort sizes and aggregate event structure (1,783 patients and 441 observed events). Synthetic data included latent composition, a composition-independent axis component, four observed composition comparators, cohort shifts, event times, and scenario-calibrated censoring. The analyzed axis was standardized within cohort, and all methods estimated a log hazard ratio per one original-axis standard-deviation unit. Endpoint-noise scenarios used balanced within-cohort label swaps so that event counts remained fixed while the observed endpoint differed from the clean biological endpoint.

Thirteen prespecified scenarios represented ideal null and signal settings, cohort shifts, noisy comparators, non-linear composition, and 5% or 10% endpoint noise. Six methods were compared: original-axis univariable Cox regression, multivariable axis-plus-comparator Cox regression, within-cohort linear residualization, principal-component residualization, and study-level cross-fitted linear or ridge residualization. Each scenario–method combination used 500 final repetitions. Cohort estimates were pooled by REML with Knapp–Hartung uncertainty; failed cohort and pooled fits were retained in the denominators. Large reference simulations defined clean-biological and observed-endpoint oracle targets. Prespecified summaries included bias, root-mean-square error, 95% interval coverage, null rejection, adverse-direction power, and fit-failure fractions.

### Module ablation and matched random-set controls

Each broad module was scored as the unweighted mean of its gene-wise z-scores and evaluated with the same cohort-specific Cox and meta-analysis framework as the full score. In the module-ablation analysis, all genes assigned to one of the five biological modules were removed before the score was recalculated. This procedure tests gene-category contribution and is distinct from leaving out a patient cohort. We also re-calculated the score after removing each gene individually.

For negative controls, 1,000 non-axis 87-gene sets were sampled with a fixed seed from genes that had finite, nonzero variance in every principal GEO cohort. Control genes were matched to the axis through 20 expression-quantile bins. The primary null statistic was the absolute pooled meta-analysis z-statistic, and the empirical value was (1 + *N*_extreme_)/(1 + 1000).

### Single-cell projection into GSE178341

The processed GSE178341 object contained 370,115 retained cells and 43,113 genes from the published CRC atlas [9]. Existing broad and detailed annotations and t-SNE coordinates were retained. All genes in the secondary 84-gene panel were present. Expression was represented as log (1 + counts per 10,000), and each panel gene was standardized across cells. The 84-gene localization score for cell *i* was

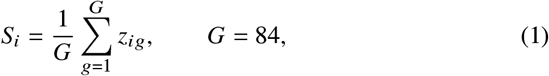

where *z*_*ig*_ is standardized expression of gene *g*. A secondary bulk-effect-weighted score was

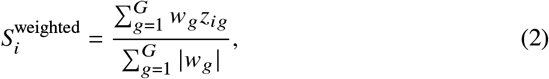

where *w*_*g*_ is the GEO `limma` log fold change. Cell-type summaries included counts, means, medians, interquartile ranges, and the proportion above the all-cell 90th percentile. These are descriptive cell-level summaries.

### Patient-level single-cell pseudobulk analysis

The patient-level analysis used the official GSE178341 UMI HDF5 matrix, metadata, and cluster table and the GEO-processed GSE132465 UMI matrix and cell annotation [9,10]. The latter is the processed count matrix distributed by GEO because original raw sequence data were not provided. A repository manifest froze the five direct GEO URLs, expected byte counts, and MD5 values; restoration aborted on any mismatch. Cells were standardized to dataset, patient, sample, tissue, and broad cell type, then counts were summed separately for every patient–tissue–broad-cell-type combination. Exact one-to-one barcode matching and absence of duplicate metadata rows were required before analysis. GSE132465’s source annotation contained one broad stromal category, whereas GSE178341 permitted fibroblast/CAF and other-stromal categories. The corrected generic mapping tested for the string `mast` before T-cell substrings; an assertion required every mast label to map to `mast_cell` and none to `T_NK_ILC`. The two datasets were analyzed independently without integration or cross-study batch correction.

The main analysis required at least 50 cells and positive total UMI count per pseudobulk profile; 20- and 100-cell thresholds were sensitivity analyses. Within each dataset and threshold, edgeR library-size normalization and trimmed mean of M-values factors were applied, followed by log counts per million with a prior count of one [21]. Each gene in the 84-gene panel was standardized across eligible profiles, multiplied by its pre-defined direction, and averaged to obtain the 84-gene signed state score. The frozen 87-gene primary bulk score was not redefined or tested in this analysis.

Tumor–normal differences were tested within broad cell type. A paired Wilcoxon test was used when at least three patients had both tissues. When fewer than three pairs remained but at least five tumor and five normal patients were available, an unpaired Wilcoxon test was reported as exploratory; otherwise the comparison was marked non-estimable. Benjamini–Hochberg adjustment was performed across broad cell types within dataset, threshold, and score. Cell-type abundance was the cell-type count divided by all retained cells from that patient and tissue and was tested analogously. Leave-one-patient-out analysis was performed for the full signed score at the 50-cell threshold.

### Spatial-transcriptomic projection

GSE267401 comprises four FFPE CRC sections profiled by Visium CytAssist: primary tumors and matched liver metastases from two patients [11]. Sections were processed separately. Spots were required to be within the tissue mask, to exceed the slide-specific fifth percentiles for total counts and detected genes, and to have mitochondrial fractions no greater than 25%. A first-percentile count and gene threshold was used as a sensitivity analysis. The resulting spot counts were normalized as log (1 + 10,000 × count / library size).

For the primary spatial projection, each available frozen-score gene was standardized across spots within its section and the 87 standardized values were averaged without weights. The 84-gene signed state score was used as a sensitivity analysis. Exploratory CAF and stromal proxies contained 11 and 14 genes, respectively; every proxy gene was also a frozen-score member. Within-section Spearman correlations quantified score agreement and association with compartment proxies. Spatial adjacency was constructed from the Visium coordinates. Moran’s *I* and neighborhood tests used 1,000 permutations, and neighbor-smoothed score agreement and upper-quartile spatial overlap were also summarized.

For the gene-disjoint sensitivity analysis, externally curated candidate CAF and stromal/perivascular markers were audited against both frozen gene lists before examining spatial associations. Every overlapping marker was removed, and remaining genes had to be detected in at least 5% of spots in at least three sections. The resulting 16-gene CAF and 15-gene stromal proxies were scored by within-section gene standardization and unweighted averaging. We report section-wise Spearman correlations and descriptive patient medians. Sections, not spots, defined the available biological replication; no patient-level hypothesis test was performed with two patients. An attempted block-permutation procedure did not provide an accepted exchangeability-preserving null and was withdrawn; its *p*-value fields are retained as NA and are not used in this manuscript.

### Focused ligand–receptor expression screen

The ligand–receptor screen used the patient-level pseudobulk profiles at the 50-cell threshold and the 1,939 interactions loaded from CellChatDB.human [12]. Other stromal cells were prespecified as the sender and epithelial cells as the receiver; this shared broad pair was used because GSE132465 lacked a fibroblast/CAF-specific annotation. GSE178341 contributed five eligible sender patients and 62 receiver patients, whereas GSE132465 contributed 14 and 22, respectively. Only genes available in the 84-gene signed state resource could contribute expression evidence.

Within each dataset, expression was represented as log (1 + CPM). The screening score multiplied mean sender-ligand expression by mean receiver-receptor expression and by the square root of the two patient-level detection fractions. A pair required nonzero patient-level expression support in both datasets to pass the expression screen. This calculation did not run the CellChat probability model, did not produce communication *p* values or FDR estimates, and did not constitute communication inference. A frozen evidence table additionally required direct spatial support, outcome evidence, and reviewed prior evidence before selecting a primary pair. Multi-subunit complexes were audited consistently at the evidence-review stage; the inconsistently handled *SEMA3C*–*NRP1_NRP2* row was excluded.

### Focused EMT sensitivity analyses

Single-cell review motivated a seven-gene EMT-related sensitivity axis comprising *ZEB1, ZEB2, VIM, MMP2, COL1A1, COL3A1*, and *TAGLN*. A regulator-removal analysis repeated clustering and survival comparisons after removing *ZEB1* and *ZEB2*. These post hoc narrowing analyses are reported only in the Supplementary Information and are not used to define the principal frozen 87-gene result.

### Statistical analysis and software

Unless stated otherwise, statistical tests were two-sided. We used Cox proportional-hazards models for time-to-event analyses, Kaplan–Meier curves and log-rank tests for exploratory group comparisons, Wilcoxon tests for nonparametric comparisons, and Spearman coefficients for rank correlations. The Benjamini–Hochberg procedure controlled the false discovery rate within the specified analysis families. Effect estimates are reported with 95% confidence intervals.

Analyses were performed in R and Python. The main R packages were `survival`, `metafor`, `limma`, and `edgeR`, with EPIC and CMScaller used for tumor-microenvironment and CMS analyses. Python analyses used `numpy`, `pandas`, `scipy`, and `scikit-learn` for data processing, clustering, simulation, and spatial analyses. Package versions, random seeds, and session information are recorded in the analysis repository.

### Ethics statement

This study used only public, de-identified data and involved no participant recruitment, intervention, or access to identifiable information. The investigators of the source studies reported their ethical approvals and consent procedures in the corresponding publications and data records.

### Use of generative artificial intelligence

OpenAI Codex was used during manuscript preparation to assist with language editing, structural reorganization, LaTeX formatting, and consistency checks. It was not used to generate or alter study data or statistical results. The authors reviewed the resulting text and remain fully responsible for the accuracy, interpretation, citations, and integrity of the submission.

### Data availability

All primary data are public. GEO microarray accessions are GSE143985, GSE17536, GSE17537, GSE17538, GSE33113, GSE38832, GSE14333, GSE161158, GSE29621, GSE37892, GSE39582, and GSE72970 (https://www.ncbi.nlm.nih.gov/geo/). GSE178341 and GSE132465 provided the single-cell data, and GSE267401 provided the spatial-transcriptomic data. TCGA COAD/READ expression and clinical data are available through UCSC Xena (https://xenabrowser.net/datapages/). E-MTAB-12862 expression and sample metadata are available through BioStudies (https://www.ebi.ac.uk/biostudies/arrayexpress/studies/E-MTAB-12862); its public clinical supplement accompanies the source publication [4].

The repository contains the frozen score definition and model metadata, the secondary annotation table, endpoint and cohort maps, compact statistical outputs, simulation summaries, provenance records, automated checks, and analysis code. Large public single-cell and spatial matrices are not duplicated in the repository. The single-cell source files can be restored from frozen GEO URLs and checked against recorded byte counts and MD5 values. Manuscript-facing outputs have a SHA-256 manifest. The gene-disjoint spatial analysis is descriptive, and the withdrawn block-permutation *p*-values are not used. The core analysis package is available at GitHub commit `6c34baf` on branch `Phase_9_Submission_Enhanced`. The exploratory E-MTAB-12862 LASSO analysis is available at GitHub commit `74a9a51` on branch `Phase_9_Submission_Package`.

## Results

### Cohort assembly and analysis flow

The 12 screened GEO series contained 2,223 GPL570 samples. Progression-related follow-up was complete for 1,983 samples, including 496 events. GSE17538 is a SuperSeries that repeated 145 endpoint-complete records from GSE17536 and 55 from GSE17537. We retained the two component cohorts and removed all 200 repeated GSE17538 records, including 55 event records. No patient identifier was duplicated in the resulting analysis table. The primary cohort-level models therefore included 1,783 tumors and 441 events from 11 non-overlapping cohorts. The selected six-versus-six partition contained 472 modeled samples in the discovery arm and 1,311 in the internal assessment arm. TCGA COAD/READ contributed 376 primary tumors with 102 PFI events. E-MTAB-12862 contained 1,063 tumor profiles; 948 stage I–III patients with 436 events entered the primary RFS model, and 1,062 patients with 502 deaths entered the secondary OS model (Table 1). Figure 1 summarizes the analysis design and the separate roles of the primary and secondary gene definitions.

### The outcome-informed GEO split is descriptive

PCA of the evidence-gene matrix showed that the first component explained 31.7% of the variance and separated much of the two-cluster expression structure (Supplementary Fig. S2a,b). In the selected discovery arm, 143 tumors were assigned to the high-axis group and 329 to the low-axis group. The high-axis group had shorter PFS-like survival (log-rank *p* = 8.66 × 10^−6^; Supplementary Fig. S2c). The internal assessment arm contained 456 high-axis and 855 low-axis tumors and showed the same direction (log-rank *p* = 3.67 × 10^−6^; Supplementary Fig. S2d). These small *p* values cannot be interpreted as independent evidence because survival behavior contributed to selecting the six-versus-six partition. They describe the expression and survival pattern that motivated the later analyses.

### Survival associations differ across bulk cohorts

The primary GEO survival analysis used the frozen 87-gene continuous score rather than the groups from the outcome-informed six-versus-six partition. Higher scores were associated with shorter survival in most cohorts, although estimates from cohorts with few events were imprecise. Random-effects meta-analysis across 11 cohorts (1,783 patients; 441 events) gave a pooled hazard ratio of 1.33 per within-cohort standard-deviation increase (95% confidence interval 1.19–1.49; *p* = 0.000179; *I*^2^ = 14.4%). Leave-one-cohort-out estimates ranged from 1.31 to 1.38. In a separate leave-one-dataset-out analysis, the expression classifier was trained without the held-out cohort before group assignment; these estimates were heterogeneous and were not used as the primary evidence.

The 11 cohorts did not all measure an identical endpoint: seven used disease-free survival (866 patients; 197 events), and one cohort each used recurrence-free, metastasis-free, metastasis/recurrence-free, or strict progression-free survival. The DFS-only pooled hazard ratio was 1.39 (95% confidence interval 1.19–1.62). Excluding the single RFS, MFS, or metastasis/recurrence-free cohort in turn gave pooled hazard ratios of 1.38, 1.33, and 1.32, respectively, without a direction reversal. No single non-DFS endpoint family accounted for the pooled direction. The combined outcome nevertheless remains a progression-related composite rather than strict PFS.

The score was next evaluated in TCGA COAD/READ. Among 376 primary tumors with PFI data and 102 events, the continuous score was associated with shorter PFI (hazard ratio 1.25, 95% confidence interval 1.03–1.52; *p* = 0.0266). The frozen high-axis versus low-axis comparison gave a hazard ratio of 1.73 (1.17–2.57; *p* = 0.00598). Both exposures were calculated before the TCGA outcomes were joined.

By contrast, the frozen equal-weight score was not associated with RFS in E-MTAB-12862. All 87 genes were measured, and the score was calculated before the clinical outcomes were joined. Among 948 stage I–III patients with 436 events, the hazard ratio per score standard deviation was 1.05 (95% confidence interval 0.95–1.16; *p* = 0.346). Adjustment for categorical stage gave a hazard ratio of 1.03 (0.93–1.14; *p* = 0.525). The axis term met the proportional-hazards assumption, although stage III did not in the adjusted model. In the secondary OS analysis of 1,062 patients with 502 deaths, the hazard ratio was 1.06 (0.97–1.16; *p* = 0.193). The confidence intervals included the null and did not support effects of the magnitude estimated in GEO or TCGA (Fig. 2).

**Figure 2.**
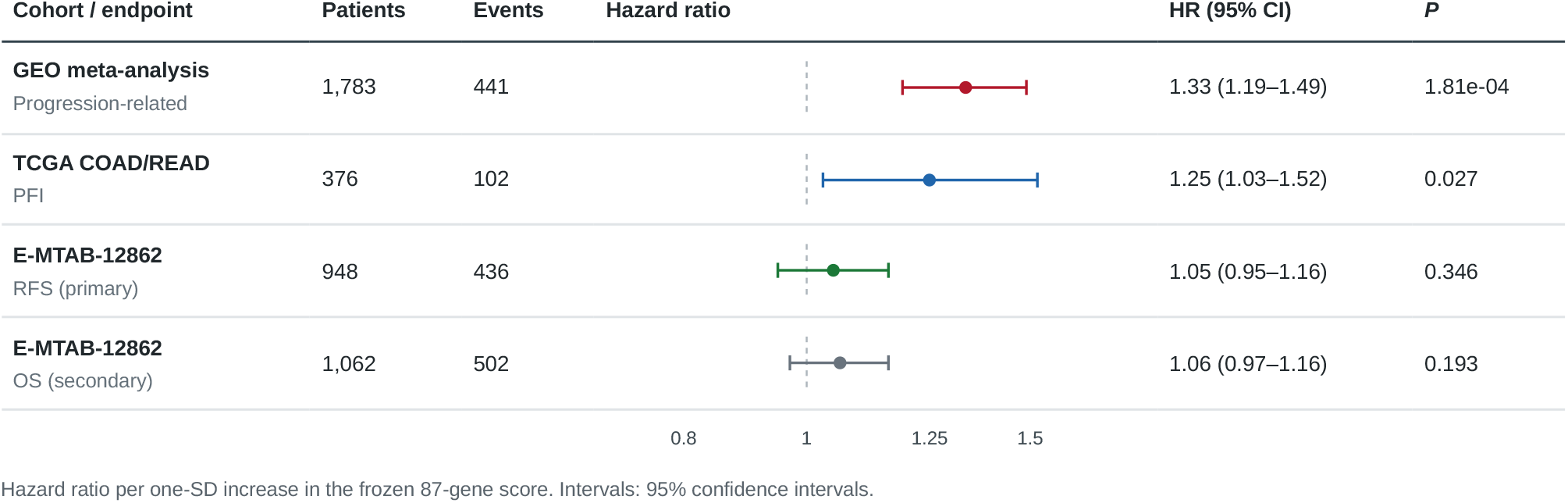
Continuous-score estimates across bulk cohorts. Hazard ratios are reported per one-standard-deviation increase in the frozen 87-gene score. GEO is a random-effects meta-analysis of 11 cohort-specific models; TCGA is the locked PFI transport analysis; E-MTAB-12862 RFS is the primary independent external assessment; and E-MTAB-12862 OS is secondary. The endpoints are related but are not identical.

After this locked analysis, RFS outcomes from E-MTAB-12862 were used to develop an exploratory LASSO score. The full-cohort model retained 34 of the 87 genes. The largest positive coefficients were assigned to *NRP1, CXCL14, PDGFRA*, and *ENO2*; the largest negative coefficients were assigned to *THY1, SEMA6D, ROBO1*, and *CXCR4*. In the same cohort used for model fitting, the standardized score had a hazard ratio of 1.80 for RFS (95% confidence interval 1.64–1.98; *p* = 8.79 × 10^−35^) and an apparent concordance index of 0.658. The mean concordance index across five held-out folds was 0.584, but the penalty had been selected from the full E-MTAB-12862 dataset before these fold-specific fits and the estimate is not from nested cross-validation.

For the prespecified descriptive display, the highest and lowest score quartiles each contained 237 patients. The high-score quartile had 172 RFS events and the low-score quartile had 66. The apparent high-versus-low hazard ratio was 4.07 (95% confidence interval 3.06–5.42; Cox *p* = 6.21 × 10^−22^; log-rank *p* = 2.66 × 10^−25^; Supplementary Fig. S6). The same-cohort result defines a candidate model for external evaluation. Its prognostic performance must be measured after the 34 coefficients have been fixed.

### High-axis tumors show a stromal and matrix-remodeling transcriptional program

Genome-wide `limma` analysis tested 22,834 genes. At false-discovery rate (FDR) below 0.05, 14,853 genes were significant; 267 also had absolute log fold change at least 1.0. At the prespecified display threshold of FDR below 0.05 and absolute log fold change at least 0.5, 1,190 genes were differentially expressed: 1,011 were higher in high-axis tumors and 179 were higher in low-axis tumors (Fig. 3a). High-axis genes included ECM and remodeling components such as *COL1A1, COL3A1, COL5A2, COL6A2, COL6A3, SPARC, VCAN*, and *TIMP2*, together with neural-guidance-associated genes including *NRP1, NRP2, SLIT2*, and *ROBO1*. Geneset enrichment identified epithelial–mesenchymal transition, extracellular-matrix organization, collagen organization, angiogenesis, cell migration, and axon-guidance programs among the high-axis signals (Fig. 3b). A 60-gene display subset showed coordinated high-axis expression across the 1,783 modeled samples (Supplementary Fig. S3).

**Figure 3.**
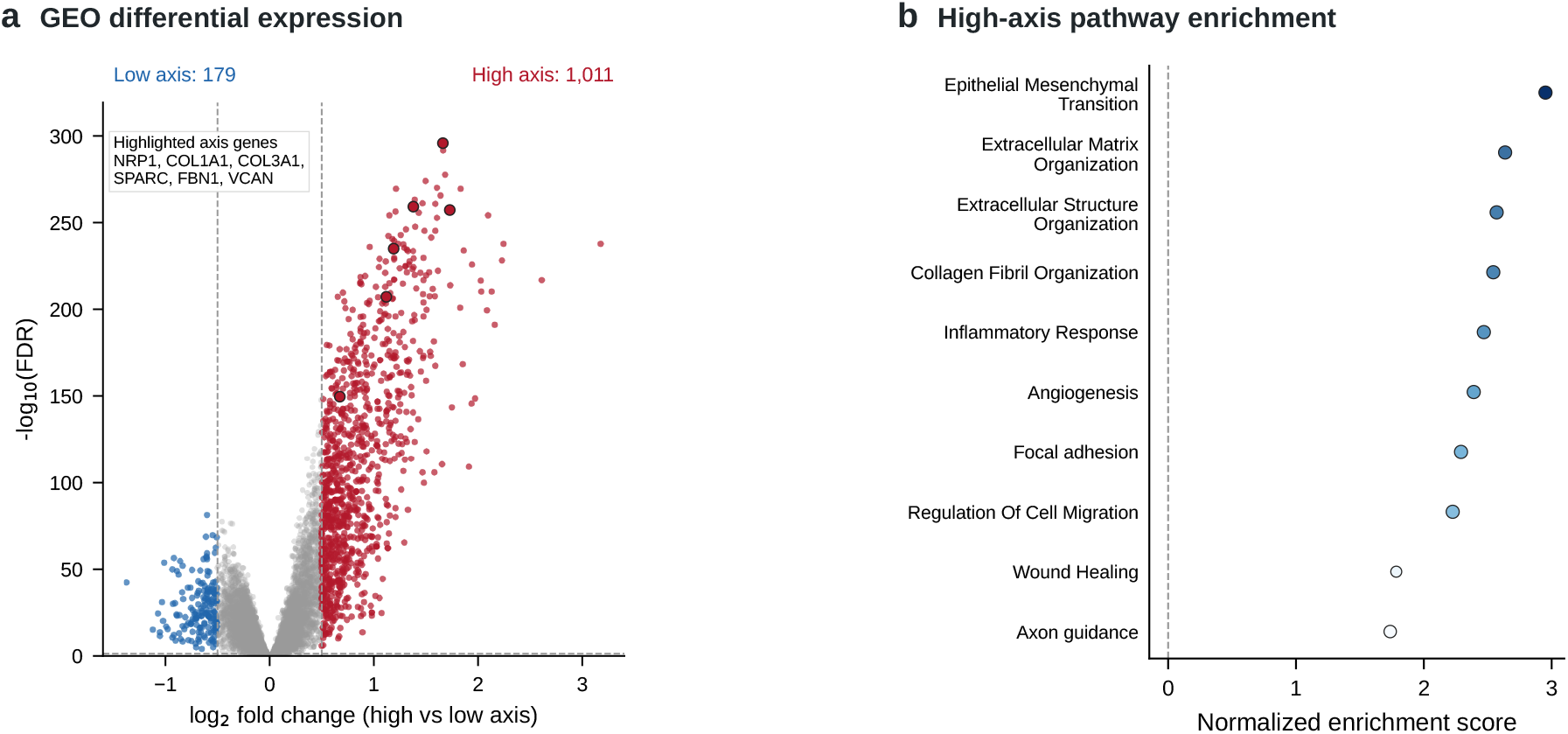
Genome-wide expression and pathway differences between GEO high- and low-axis tumors. (a) Positive log fold change denotes higher expression in the high-axis group. Colored points meet FDR < 0.05 and |log FC| ≥ 0.5; selected axis genes are outlined. (b) Selected high-axis-enriched pathways from the preranked analysis. Point size reflects −log_10_(FDR).

### Secondary module patterns and frozen-score controls implicate stromal remodeling

Dataset-standardized marker scores were concordant with the differential-expression pattern. Fibroblast, stromal, macrophage, endothelial, and immune signals were higher in high-axis tumors after adjustment for dataset, whereas the tumor-purity proxy was lower (Fig. 4a). The very small FDR values for these comparisons reflect the large pooled sample size and do not by themselves establish a clinically important difference.

**Figure 4.**
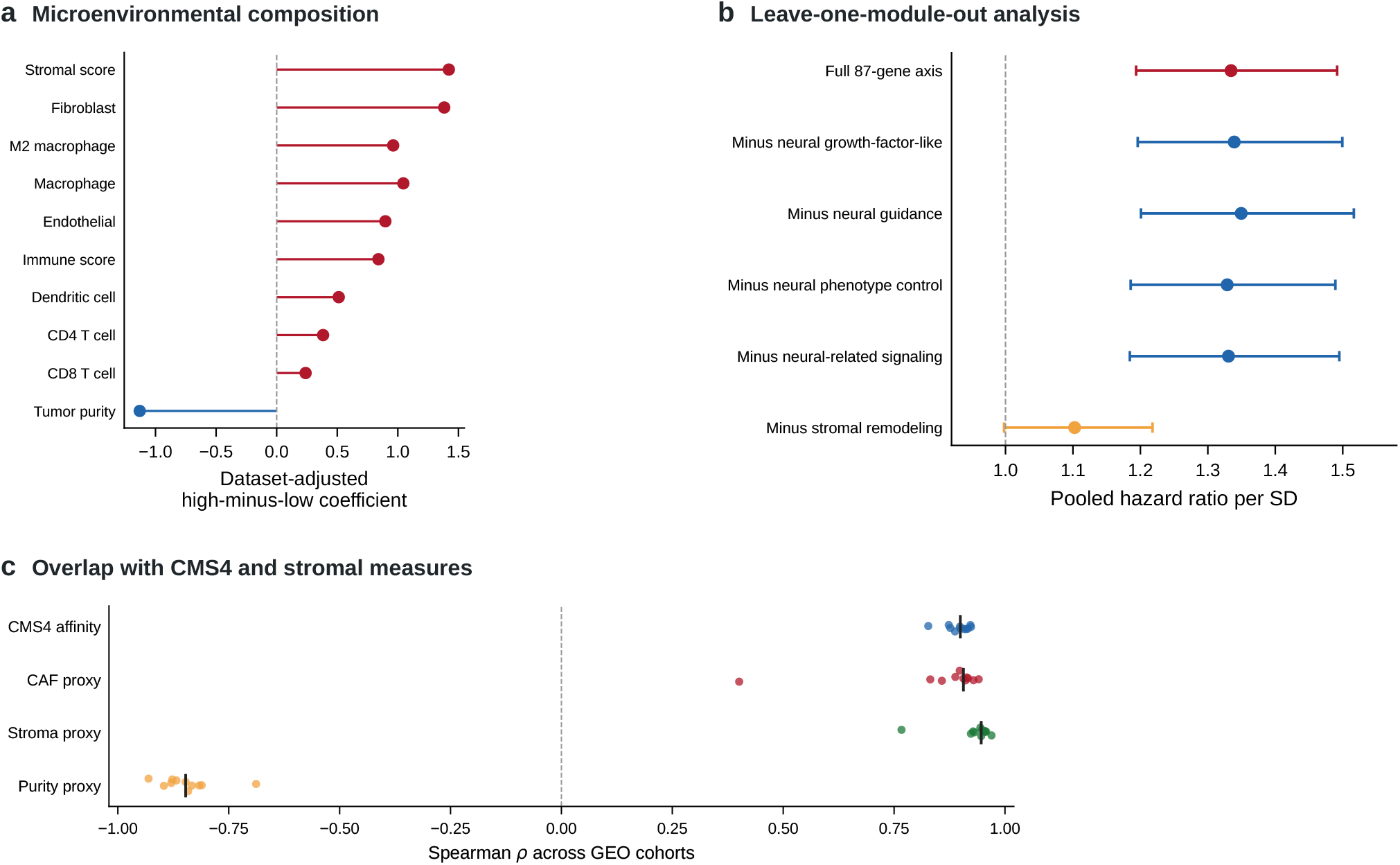
Microenvironmental composition and module controls in GEO. (a) Dataset-adjusted high-minus-low coefficients for selected dataset-standardized marker scores. Positive values indicate higher scores in high-axis tumors. (b) Random-effects estimates for the full 87-gene score and scores formed after removal of each broad module. (c) Cohort-specific Spearman correlations between the axis and CMS4 affinity, CAF, stroma, and purity proxies; black vertical marks show medians.

The stromal-remodeling module alone had a pooled hazard ratio of 1.34 (95% confidence interval 1.18–1.53), close to the full-axis estimate of 1.34. Removing all 55 stromal-remodeling genes reduced the estimate to 1.10, whereas removing all 15 neural-guidance genes left it at 1.35 (Fig. 4b). Removing any single gene did not reverse the direction or raise the meta-analysis *p* value above 0.05. Three of 1,000 expression-matched random gene sets had an absolute metaanalysis z-statistic at least as large as the observed score (empirical *p* = 0.003996). These analyses indicate that the GEO association does not depend on one gene and is accounted for mainly by the stromal-remodeling module.

### The axis captures a stromal component within the broader CMS4 phenotype

Within the GEO cohorts, the frozen score correlated with CMS4 distance-derived affinity (median Spearman ρ = 0.904), EPIC CAF or a labeled CAF proxy (ρ = 0.909), and a stromal proxy (ρ = 0.947); its correlation with the purity proxy was negative (ρ = −0.858; Fig. 4c). CMScaller template distance was converted to a continuous affinity measure and was not treated as a posterior probability. Adding the axis to models that already contained CMS4 and one stromal comparator did not improve the primary CMS-adjusted model likelihood. Several models also showed substantial collinearity. The overlap places the axis within the mesenchymal and stromal biology covered by the broad CMS4 classification.

For the cross-fitted analysis, a nuisance model learned the relationship between the axis and CMS4, CAF, stroma, and purity from ten GEO cohorts and was then applied to the held-out cohort. Each residual remained on the original-axis scale and was not re-standardized. The pooled residual-axis hazard ratio was 1.12 (95% confidence interval 0.77–1.64; prediction interval 0.76–1.65; *I*^2^ = 16.2%). A prespecified ridge model gave a similar result. Training the nuisance model in all GEO cohorts and applying it once to TCGA gave a residual-axis hazard ratio of 1.31 (0.63–2.74). The intervals were wide, but neither analysis detected an association after removal of the measured composition components.

Seven GEO cohorts with usable categorical stage contributed 943 patients and 225 events. After stage adjustment, the original score remained associated with outcome (hazard ratio 1.26, 95% confidence interval 1.06–1.50; *p* = 0.0165), but the cross-fitted residual did not (0.94, 0.62–1.41; *p* = 0.706).

E-MTAB-12862 provided a separate composition check. The score correlated with the source CMS4 binary label (ρ = 0.699), a CAF marker proxy (ρ = 0.953), a stromal proxy (ρ = −0.954), and an inverse purity proxy (ρ = −0.909). The stromal module RFS estimate was nearly identical to the full-score estimate (hazard ratios 1.05 and 1.05), whereas the neural-guidance module estimate was 0.98. Applying the GEO-trained nuisance model gave an RFS residual-axis hazard ratio of 1.001 (0.998–1.004). E-MTAB reproduced the score’s stromal composition pattern without reproducing its prognostic association. The comparator analyses did not identify additional prognostic information after accounting for CMS4 and generic stromal abundance. These results place the axis primarily within the stromal-remodeling component of CMS4 and do not support a separate neural or prognostic subtype.

### Simulation defines the boundary conditions of composition residualization

The frozen simulation benchmark crossed 13 null, signal, cohort-shift, comparator-noise, nonlinear-composition, and endpoint-noise scenarios with six analysis methods and 500 final repetitions, yielding 39,000 pooled meta-analysis fits. All pooled fits completed; 472 of 429,000 cohort-level fits failed and remained in the failure accounting. Under the ideal oracle-composition null, cross-fitted linear residualization had a false-positive rate of 0.038 and 95% interval coverage of 0.962. The false-positive rate increased to 0.102 when composition comparators were noisy and to 0.248 when the nuisance relationship was nonlinear. With 10% endpoint-label noise, the small-signal oracle log-hazard-ratio target changed from 0.161 for the clean biological endpoint to 0.131 for the observed endpoint. Cross-fitting prevents held-out-cohort leakage but does not correct measurement error or nuisance-model misspecification. The benchmark identifies these failure conditions and does not establish that the method is unbiased in patient data.

### TCGA expression patterns are consistent with the transported stromal state

The transported-group visualization is shown in Fig. 5a as descriptive support for the locked survival estimates reported above. TCGA differential expression reproduced the high-axis-skewed pattern, with extracellular-matrix and stromal genes increased in the high-score state (Fig. 5b).

**Figure 5.**
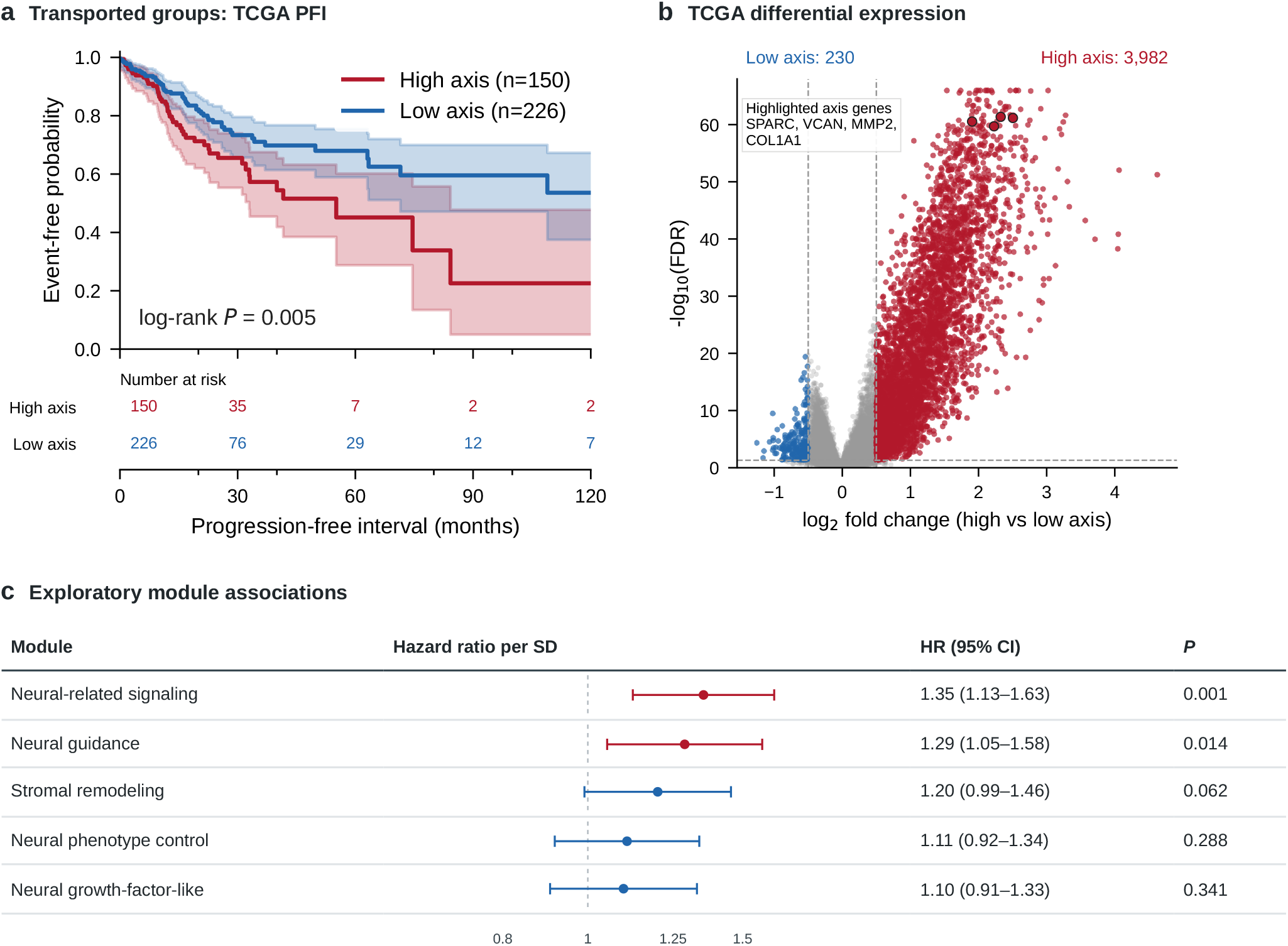
TCGA transport and secondary module analyses. (a) Kaplan–Meier PFI curves and numbers at risk for the transported GEO-derived classification; shading shows 95% confidence intervals. (b) limma differential expression; positive log fold change denotes higher expression in high-axis tumors, and selected axis genes are outlined. (c) Exploratory hazard ratios and 95% confidence intervals per standardized broad-module score. The primary locked TCGA survival estimates use the frozen 87-gene score and are reported in the text.

Exploratory Cox models for the five broad modules in the 87-gene definition produced heterogeneous estimates (Fig. 5c). These module results do not replace the locked full-score analysis or the GEO module-control analysis.

### The secondary 84-gene panel localizes to fibroblast-rich compartments

All genes in the related 84-gene panel were present in the processed GSE178341 object. Fifteen had low detection but were retained to preserve the predefined membership. The unsigned 84-gene localization score was highest in stromal regions of the published global t-SNE (Fig. 6a,b). Among broad cell types, CAFs/fibroblasts had the highest median localization score (1.827), followed by other stromal cells (0.966), endothelial cells (0.408), and macrophage/myeloid cells (0.079; Table 2). Strong gene-level contributors included *LUM, DCN, COL1A2, COL3A1, MMP2, COL6A2, COL1A1*, and *COL6A3*. Representative t-SNE expression maps for *LUM, SNAI2, ZEB2, VIM, COL1A1*, and *NRP2* are shown in Supplementary Fig. S7. This analysis locates the secondary panel across cell types; it does not redefine the frozen 87-gene primary bulk score.

**Table 2.**
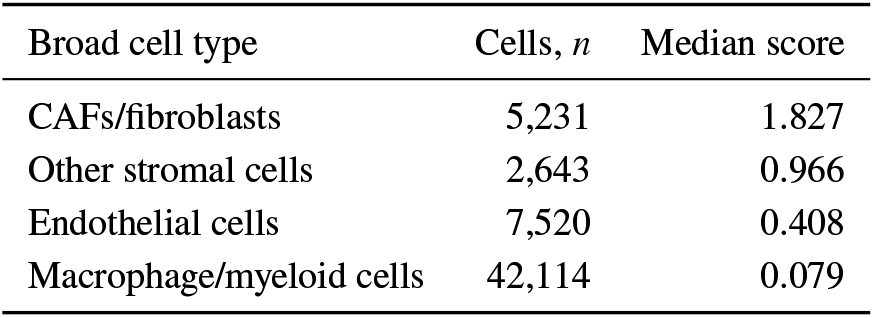
Broad cell types with the highest median 84-gene localization scores in GSE178341. Values are descriptive summaries across cells and do not represent patient-level independent observations.

| Broad cell type | Cells, *n* | Median score |
| --- | --- | --- |
| CAFs/fibroblasts | 5,231 | 1.827 |
| Other stromal cells | 2,643 | 0.966 |
| Endothelial cells | 7,520 | 0.408 |
| Macrophage/myeloid cells | 42,114 | 0.079 |

**Figure 6.**
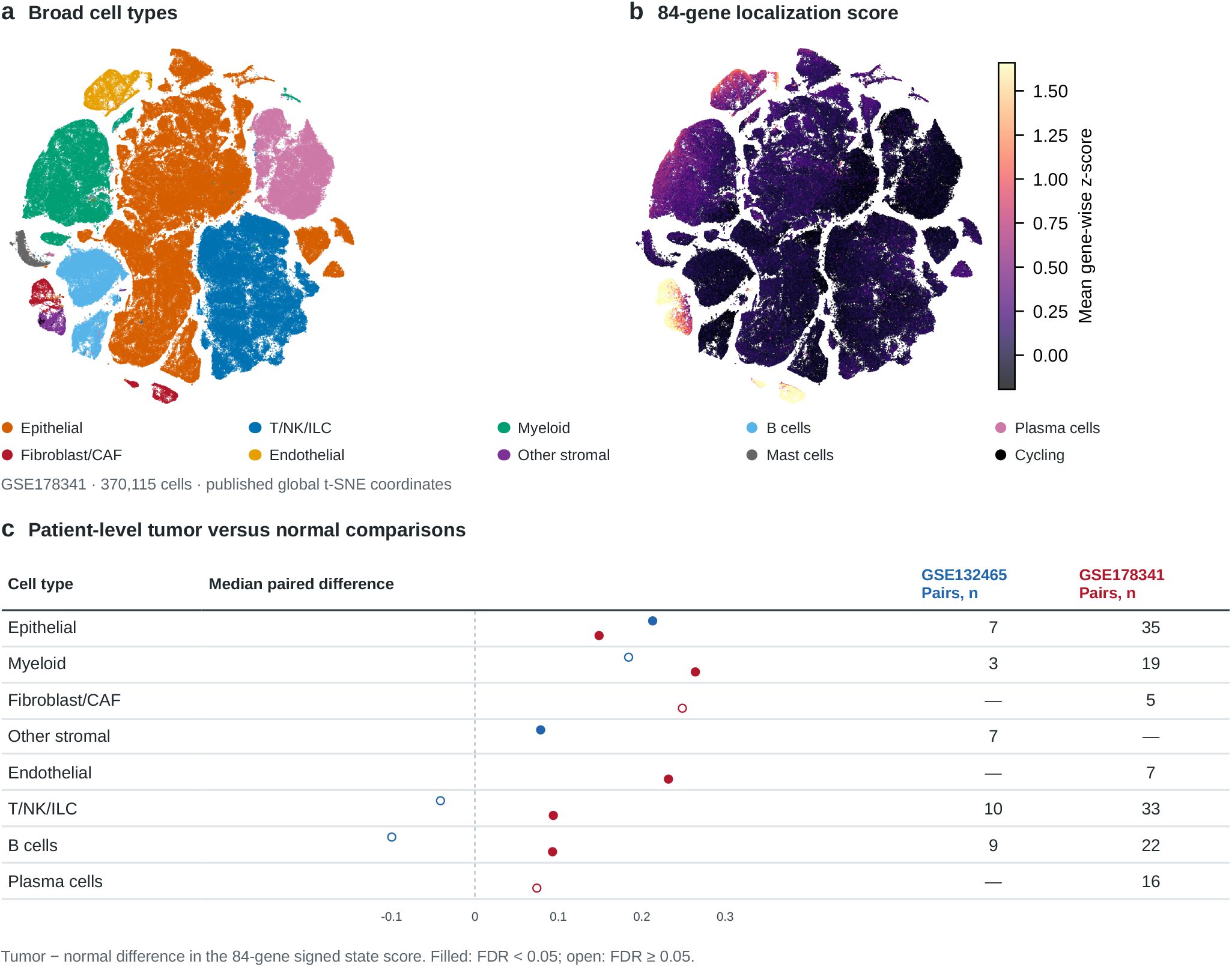
Cellular localization and patient-level single-cell comparisons. (a) Published GSE178341 global t-SNE coordinates colored by broad cell type, with a separate color legend. The Cycling group contains 44 proliferating pericytes that were kept separate from other stromal cells. (b) The unsigned 84-gene localization score on the same coordinates; the color scale is capped at the first and 99th percentiles. Panels a and b display all 370,115 cells descriptively. (c) Median paired tumor-minus-normal differences in the 84-gene signed state score at the 50-cell threshold. Blue and red points denote GSE132465 and GSE178341, respectively. The right-hand columns report paired-patient counts; a dash indicates that no completed paired comparison was available. Filled points have Benjamini–Hochberg FDR below 0.05, and open points do not. Patient is the inferential unit in panel c.

To avoid treating cells as independent replicates, we next aggregated raw or GEO-processed UMI counts into patient– tissue–broad-cell-type pseudobulk profiles. GSE178341 contributed 370,115 cells from 62 patients (64 tumor and 36 normal samples), and GSE132465 contributed 63,689 cells from 23 patients (23 tumor and ten matched normal samples). A clean rerun restored five official GEO supplementary files from a frozen URL, byte-count, and MD5 manifest; every matrix barcode matched the official metadata in both datasets. Both datasets detected all 84 annotation genes and were analyzed separately without cross-dataset integration or joint batch correction.

The corrected mapping assigned all 187 GSE132465 cells labeled as mast cells to a separate mast-cell compartment and none to T/NK/ILC. At the main minimum of 50 cells per pseudobulk profile, the 84-gene signed state score was higher in tumor in epithelial profiles (GSE132465 median paired difference 0.213, seven pairs, *p* = 0.0156, FDR 0.0391; GSE178341 0.149, 35 pairs, *p* = 1.47 × 10^−8^, FDR 1.33 × 10^−7^) and myeloid profiles (0.184, three pairs, *p* = 0.25; 0.264, 19 pairs, *p* = 3.81 × 10^−6^, FDR 1.14 × 10^−5^). The GSE132465 other-stromal score was also tumor-higher (0.079, seven pairs, *p* = 0.0156, FDR 0.0391); GSE178341 other-stromal profiles showed the same direction in an explicitly exploratory unpaired comparison (median difference 0.283, *p* = 0.00311) because only two matched pairs remained. GSE132465 provided only a broad stromal label and therefore could not independently resolve fibroblasts/CAFs. In GSE178341, the fibroblast/CAF score was tumor-higher (median paired difference 0.249, five pairs, *p* = 0.0625, FDR 0.0776) and retained the same direction at 20-, 50-, and 100-cell thresholds (Fig. 6c).

The abundance analysis did not support simple expansion of the high-scoring stromal compartments. Fibroblast/CAF proportion was lower in GSE178341 tumors than matched normal samples (32 pairs, *p* = 0.000398), while the within-fibroblast score was tumor-higher. Similarly, GSE132465 other-stromal abundance was normal-higher (ten pairs, *p* = 0.00977) despite a tumor-higher state score. Leave-one-patient-out analyses at the 50-cell threshold produced no completed direction reversal; three GSE132465 myeloid omissions reduced the matched set from three to two and made that test non-estimable. These findings support a tumor-associated state change within multiple compartments, including stroma, but do not establish a cell-intrinsic mechanism or single-cell prognostic association.

### Spatial projection is concordant with CAF- and stroma-rich regions in four sections

The frozen 87-gene score was projected onto four GSE267401 Visium CytAssist sections: primary CRC and matched liver metastases from patients 17 and 21. After slide-specific quality control, 16,171 spots were retained (3,534–4,516 per section), and all 87 score features were available in every section. H&E reference images and section-level score maps are shown in Supplementary Fig. S8. The spot-level score correlated positively with the 11-gene CAF proxy in all four sections (Spearman ρ = 0.651–0.895) and with the 14-gene stromal proxy (ρ = 0.681–0.915). Moran’s *I* ranged from 0.242 to 0.730, with empirical permutation *p* values of approximately 0.001, indicating non-random spatial organization. An 84-gene signed state score used as a sensitivity analysis was concordant with the frozen projection (ρ = 0.745–0.942), and the principal directions were unchanged under a less stringent spot-quality threshold.

These associations are descriptive rather than independent validation: the spatial sections represent only two patients, each spot may contain multiple cell types, and all genes in both original CAF and stromal proxies are members of the 87-gene score. We therefore repeated the comparison after removing every candidate marker present in either the 87- or 84-gene list and applying a prespecified detection filter. The final disjoint CAF and stromal proxies contained 16 and 15 genes, respectively, with zero overlap with either axis. Correlations remained positive in all four sections (CAF ρ = 0.227–0.814; stroma ρ = 0.373–0.808). Median section correlations were positive for both patients, but with only two patients these summaries cannot establish patient-level replication or progression association. Neural-guidance and stromal scores were only weakly correlated in one section (ρ = 0.077) and moderately correlated in the others (ρ = 0.316–0.488), providing no consistent spatial evidence for an autonomous neural component.

### A ligand–receptor expression screen does not nominate a primary interaction

A prespecified other-stromal-to-epithelial screen used patient-level pseudobulk profiles from GSE178341 and GSE132465. The broad other-stromal sender was selected because GSE132465 did not provide a fibroblast/CAF-specific label. Among 1,939 interactions loaded from CellChatDB.human, 12 pairs had nonzero patient-level ligand expression in the sender and receptor expression in the receiver in both datasets. These included *CXCL12*–*CXCR4*/*ACKR3, IGF1*–*IGF1R*, and *NGF*–*NGFR*/*NTRK1*. An initially retained *SEMA3C*– *NRP1_NRP2* complex was excluded because receptor-subunit handling was inconsistent between ranking and patient-level evidence review.

No pair met the combined requirements for replicated expression, direct spatial support, outcome evidence, and reviewed prior evidence. Direct ligand–receptor spatial testing was not performed, and the screen did not estimate CellChat communication probabilities or significance. Accordingly, no primary interaction was selected and these results are interpreted as a negative expression screen, not as evidence of intercellular communication.

### A single-gene control does not reproduce the multigene association

Clustering 1,783 GEO tumors by *VIM* expression alone produced 819 VIM-high and 964 VIM-low samples but no PFS-like difference (log-rank *p* = 0.313). The univariable group comparison (hazard ratio 1.10, *p* = 0.328) and the continuous expression model (hazard ratio 0.98, *p* = 0.718) were also null (Supplementary Fig. S5). In E-MTAB-12862, standardized *VIM* expression gave an RFS hazard ratio of 1.06 (0.96– 1.18; *p* = 0.239). The multigene GEO association cannot be attributed to *VIM* alone.

## Discussion

The frozen 87-gene score was associated with shorter progression-related survival in the GEO meta-analysis and with shorter PFI after locked transport to TCGA. The association did not replicate in E-MTAB-12862, which contributed 436 RFS events compared with 441 progression-related events across the 11 GEO cohorts. Its RFS and OS estimates were close to the null, and stage adjustment did not reveal an association. The E-MTAB result limits the generalizability of the earlier prognostic estimates. Analyses of composition and cellular localization gave a more consistent biological interpretation of the score.

CMS4 is a broad mesenchymal subtype that encompasses stromal infiltration, matrix remodeling, angiogenesis, and TGF-β-related programs [5, 6]. The score correlated closely with CMS4, CAF, and stromal measures in GEO and E-MTAB-12862. In GEO, the 55-gene stromal-remodeling module produced an estimate similar to the full score, and removing that module reduced the association. Removing the neural-guidance module had little effect. Composition residualization did not detect an additional association in GEO, TCGA, or E-MTAB-12862, although the GEO and TCGA intervals were wide. Together, these results indicate that the axis mainly reflects fibroblast-rich matrix remodeling within CMS4. They did not support prognostic information beyond the measured CMS4 and stromal components.

The single-cell and spatial results were consistent with this interpretation. The unsigned 84-gene localization score was highest in CAF/fibroblast and other stromal compartments. Patient-level comparisons with the 84-gene signed state score detected tumor-associated increases within epithelial, myeloid, and stromal profiles where those comparisons were estimable. The spatial score formed organized regions and correlated positively with CAF and stromal proxies even after genes shared with the 87- and 84-gene lists were removed from those proxies. These analyses locate the expression program in fibroblast-rich tissue regions and tumor-associated cellular states. They do not test patient prognosis because the single-cell datasets lacked outcome data and the spatial dataset contained only two patients.

The module analyses also refine the meaning of the neural label. Neural-guidance genes contributed little to the GEO ablation result, and the neural-guidance module was not associated with RFS in E-MTAB-12862. Axon-guidance genes can participate in cell migration, vascular patterning, and stromal signaling without indicating neuronal differentiation. In this study, neural guidance is therefore a gene-category label, not evidence of a tumor-cell-intrinsic neural mechanism. The ligand–receptor screen identified 12 pairs with minimal sender and receiver expression support in both single-cell datasets, but none met the full evidence criteria. Co-expression alone does not establish physical proximity, signaling activity, direction, or an effect on progression.

The E-MTAB result changes the interpretation of the earlier survival findings. Platform, endpoint, stage distribution, treatment, and cohort composition differ among GEO, TCGA, and E-MTAB-12862, but the present analyses cannot determine which difference explains the attenuation. The exploratory 34-gene LASSO model shows that outcome-derived weights can improve separation within E-MTAB-12862: its apparent concordance index was 0.658, compared with a mean of 0.584 in held-out folds that used a penalty selected from the full cohort. These estimates describe model development within EMTAB-12862. The analysis defines a candidate 34-gene score. Its weights should be fixed before another cohort is used to estimate prognostic performance.

The study has additional limitations. The original sixversus-six GEO partition was selected through a search that included survival behavior, so its Kaplan–Meier results are exploratory. The cohort-level meta-analysis avoids treating the selected split as validation, but it remains a retrospective analysis of heterogeneous endpoints. Seven cohorts measured DFS, and each other endpoint family was represented by one cohort. The umbrella term PFS-like does not make these outcomes equivalent. Clinical covariates were incomplete in GEO, several cohorts had few events, and the stage-adjusted analysis included only seven cohorts. The E-MTAB stage-adjusted model also showed non-proportionality for stage III.

The 87-gene list was assembled from bladder-cancer and broader cancer literature rather than a CRC-specific systematic review. The primary 87-gene score and the secondary 84-gene localization and signed state scores have different membership and scoring rules and cannot be treated as interchangeable versions of one model. Composition proxies are also imperfect. The simulation showed that cross-fitting prevents outcome leakage into residual construction, but noisy measurements and nonlinear nuisance relationships can still produce biased inference. Gene-disjoint spatial proxies remove direct gene reuse but do not remove shared tissue architecture or other sources of composition confounding.

The survival associations differed by cohort, but the biological analyses converged on a consistent interpretation. Across bulk, single-cell, and spatial data, the axis mainly reflected a fibroblast-rich, matrix-remodeling state within the broader CMS4 phenotype. This result links the initial outcome association to a defined tissue program and provides a testable hypothesis about stromal organization in CRC progression. Current evidence does not support use as a standalone prognostic biomarker. Future prognostic studies should fix the scoring rule in advance, use a uniform clinical endpoint, and evaluate it in a cohort that was not used for model development. Functional experiments are needed to test the cellular mechanisms represented by the score.

## Supporting information

Supplemental Results

## Acknowledgements

None.

## Author contributions

Y.Z. (Tia) performed the main research work, including data curation, formal analysis, investigation, visualization, and manuscript drafting. S.H. (Shuqiang Hao) and C.J. (Chonghe Jiang) contributed to the study concept, discussed and refined the ideas, and supervised the work. All authors interpreted the results, contributed to manuscript review, and approved the final manuscript.

## Additional information

Supplementary Information accompanies this paper.

