## Supplemental Results for "A fibroblast-rich transcriptomic axis reflects stromal remodeling within the CMS4 phenotype in colorectal cancer"

### Supplementary Methods and Results

#### Frozen gene definitions and provenance

The authoritative primary score contains 87 genes, all with positive direction, and is calculated as the unweighted mean of within-cohort gene-wise z-scores. The source evidence table was assembled before the present CRC outcome analyses from bladder-cancer and broader cancer literature. It is therefore a cross-cancer biological hypothesis and not a CRC-specific literature signature. The outcome analyses reported here did not revise the frozen membership or direction.

The later 84-gene panel is a secondary annotation resource. It contains 55 stromal-remodeling genes, 15 neural-guidance genes, eight neural-related phenotype-control genes, five neural-related signaling genes, and an EGFR special-check gene. It shares 83 genes with the primary list, excludes *ERBB2*, *ERBB3*, *ERBB4*, and *NRG1*, and adds *EGFR*. The local audit confirmed that the four excluded genes were measured in the principal GEO cohorts, so the membership change was not a response to missing bulk expression. The source files do not document why the later panel was changed. We used its membership without directions for the descriptive 84-gene localization score and used its predefined directions for the patient-level 84-gene signed state score. Neither score redefines the frozen 87-gene primary score. The complete definitions, comparison, cohort manifest, and endpoint map are supplied in `supplementary_data/gene_definitions/`.

The GEO overlap audit found that the 200 endpoint-complete GPL570 records in GSE17538 repeated 145 records from GSE17536 and 55 from GSE17537. These repeated records included 55 events. We retained the two component series and excluded the corresponding GSE17538 records before cohort-level modeling. The final 11-cohort table contained 1,783 unique patient identifiers and 441 events, with no between-cohort patient duplication.

#### Post hoc Harmony and scVI representation checks

Harmony and scVI were applied after the original PFS-associated labels had been fixed. They did not discover or reassign risk groups. Both checks used 22,834 common genes and the same six-cohort discovery arm and six-cohort internal assessment arm described in the main manuscript.

For Harmony, within-dataset standardized expression matrices were combined, PCA was performed, and dataset identity was supplied as the batch variable. UMAP visualized the corrected coordinates. Dataset silhouettes near or below zero indicated little dataset-specific separation; positive but small risk-label silhouettes indicated weak retention of the original axis labels (Table S2).

For scVI, processed microarray intensities were converted to pseudo-counts by back-transformation, replacement of nonfinite values, clipping at zero, normalization to a total of 10,000, and integer rounding. Separate 10-dimensional models were trained for the two arms for 400 epochs with dataset as batch key. Because these inputs were not raw counts and did not arise from single-cell RNA sequencing, scVI results are exploratory.

The original and corrected representations retained similar sample relationships. Canonical correlations were strongest between Harmony and scVI and exceeded 0.95 for the first two canonical variates in both arms (Table S3). Canonical correlation is invariant to sign flips and rotations of individual latent dimensions, making it more appropriate here than direct axis-to-axis comparison.

#### GEO and TCGA heatmaps

The GEO heatmap displays the 60 genes with the largest absolute high-versus-low mean difference among the 84 genes available in the standardized display matrix. It is a visualization subset rather than a new score definition. The 1,783 tumors are ordered by axis group and continuous score. Annotation bars show axis group and source cohort.

**Table S1. Frozen 87-gene primary axis.** Every gene contributes with equal weight and positive direction after within-cohort standardization. Module codes are defined below.

| No. | Gene symbol | Module | No. | Gene symbol | Module |
|---|---|---|---|---|---|
| 1 | *ERBB2* | GF | 45 | *VEGFB* | SR |
| 2 | *ERBB3* | GF | 46 | *CDH1* | SR |
| 3 | *ERBB4* | GF | 47 | *CDH2* | SR |
| 4 | *NRG1* | GF | 48 | *SNAI1* | SR |
| 5 | *DCC* | NG | 49 | *SNAI2* | SR |
| 6 | *EFNA3* | NG | 50 | *TGFB1* | SR |
| 7 | *EFNB2* | NG | 51 | *TGFBR1* | SR |
| 8 | *EPHB2* | NG | 52 | *TGFBR2* | SR |
| 9 | *EPHB3* | NG | 53 | *TGFBR3* | SR |
| 10 | *EPHB4* | NG | 54 | *TWIST1* | SR |
| 11 | *NRP1* | NG | 55 | *TWIST2* | SR |
| 12 | *NRP2* | NG | 56 | *VIM* | SR |
| 13 | *ROBO1* | NG | 57 | *ZEB1* | SR |
| 14 | *ROBO3* | NG | 58 | *ZEB2* | SR |
| 15 | *SEMA3A* | NG | 59 | *COL1A1* | SR |
| 16 | *SEMA3C* | NG | 60 | *COL1A2* | SR |
| 17 | *SEMA4D* | NG | 61 | *COL3A1* | SR |
| 18 | *SEMA6D* | NG | 62 | *COL5A1* | SR |
| 19 | *SLIT2* | NG | 63 | *COL6A1* | SR |
| 20 | *ASCL1* | PC | 64 | *COL6A2* | SR |
| 21 | *CHGA* | PC | 65 | *COL6A3* | SR |
| 22 | *ENO2* | PC | 66 | *DCN* | SR |
| 23 | *INSM1* | PC | 67 | *FN1* | SR |
| 24 | *NCAM1* | PC | 68 | *LUM* | SR |
| 25 | *NEUROD1* | PC | 69 | *SPARC* | SR |
| 26 | *POU2F3* | PC | 70 | *VCAN* | SR |
| 27 | *SYP* | PC | 71 | *ISG15* | SR |
| 28 | *BDNF* | NS | 72 | *SLC14A1* | SR |
| 29 | *NTRK2* | NS | 73 | *STAT1* | SR |
| 30 | *NGF* | NS | 74 | *STAT2* | SR |
| 31 | *NTRK1* | NS | 75 | *WNT5A* | SR |
| 32 | *NGFR* | NS | 76 | *MMP11* | SR |
| 33 | *ACTA2* | SR | 77 | *MMP14* | SR |
| 34 | *PDGFRB* | SR | 78 | *MMP2* | SR |
| 35 | *POSTN* | SR | 79 | *ACKR3* | SR |
| 36 | *RGS5* | SR | 80 | *CCL2* | SR |
| 37 | *TAGLN* | SR | 81 | *CXCL1* | SR |
| 38 | *THY1* | SR | 82 | *CXCL12* | SR |
| 39 | *FAP* | SR | 83 | *CXCL14* | SR |
| 40 | *FGF2* | SR | 84 | *CXCL2* | SR |
| 41 | *FGF7* | SR | 85 | *CXCR4* | SR |
| 42 | *IGF1* | SR | 86 | *IL6* | SR |
| 43 | *IGF1R* | SR | 87 | *PDGFRA* | SR |
| 44 | *VEGFA* | SR | | | |

SR, stromal remodeling (55 genes); NG, neural-guidance signaling (15); PC, neural-related phenotype control (8); NS, neural-related signaling (5); GF, neural-growth-factor-like signaling (4). Read the left block from 1 to 44, then the right block from 45 to 87. ACKR3 is the symbol used for the CXCR7 alias.

**Table S2. Representation-check summary.** NR, not reported in the source run summary.

| Method | Arm | Samples | Genes | Dataset silhouette | Risk silhouette |
|---|---|---|---|---|---|
| Harmony | Discovery | 672 | 22,834 | −0.05218 | 0.06835 |
| Harmony | Assessment | 1,311 | 22,834 | −0.06715 | 0.08502 |
| scVI | Discovery | 672 | 22,834 | −0.02491 | 0.082 |
| scVI | Assessment | 1,311 | 22,834 | NR | NR |

**Table S3. Canonical correlations among representations.** Each cell reports the first two canonical correlations.

| Comparison | Discovery arm | Assessment arm |
|---|---|---|
| Original PCA versus Harmony | 0.957, 0.903 | 0.959, 0.917 |
| Original PCA versus scVI | 0.932, 0.826 | 0.931, 0.852 |
| Harmony versus scVI | 0.970, 0.951 | 0.969, 0.960 |

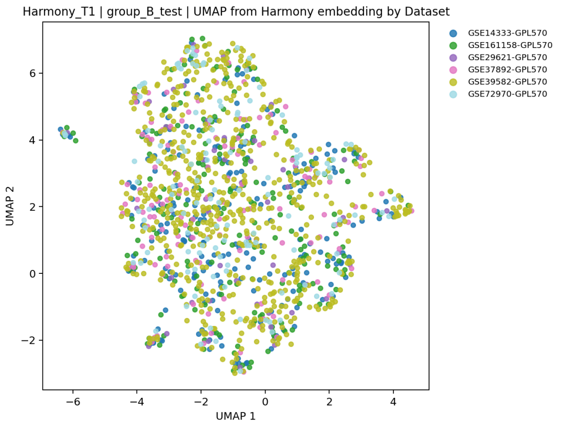


**(a)** Assessment arm, Harmony UMAP.

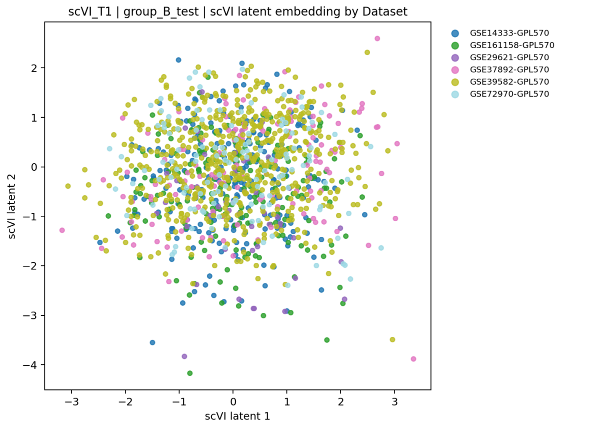


**(b)** Assessment arm, scVI coordinates.

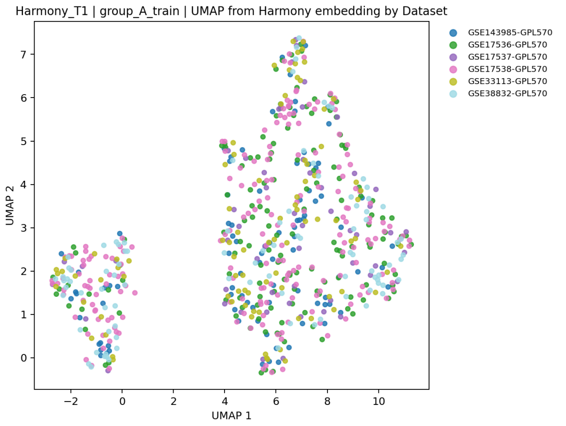


**(c)** Discovery arm, Harmony UMAP.

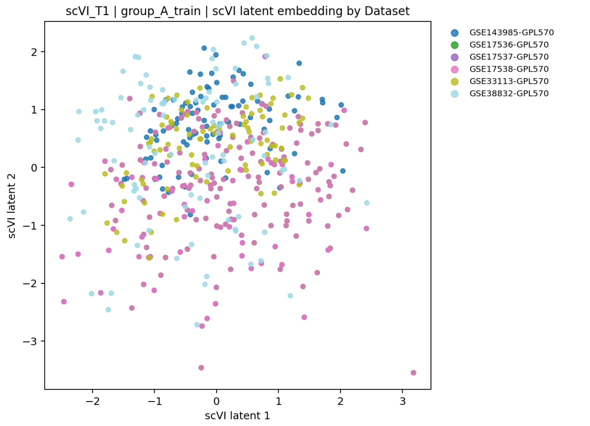


**(d)** Discovery arm, scVI coordinates.

**Figure S1. Dataset-colored representation checks.** (a,b) Internal assessment arm. (c,d) Discovery arm. The dataset labels are broadly intermingled. Harmony and scVI were fitted to the same complete-case samples within each arm. The scVI panels use pseudo-count-transformed microarray data and are exploratory. These plots assess dataset-associated structure and do not validate risk-group separation.

#### *VIM* single-gene negative control

One-dimensional $k = 2$ clustering of standardized *VIM* expression assigned 964 samples to VIM-low and 819 to VIM-high. Mean source-scale expression was 7.37 and 9.14, respectively. Despite clear expression separation, PFS-like

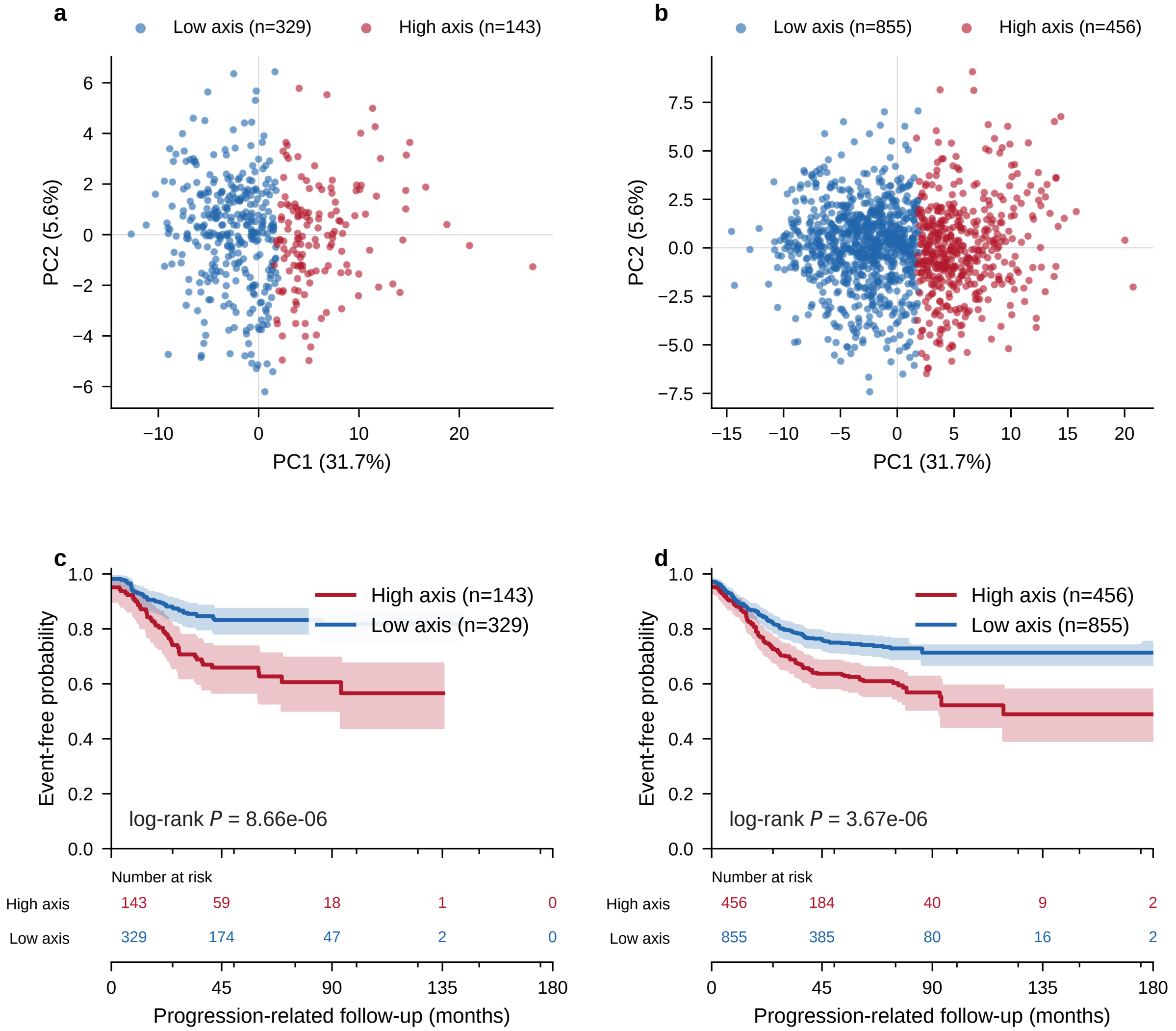


**Figure S2. Expression structure and progression-related survival in the outcome-informed GEO split.** (a,b) PCA of the evidence-gene matrix with $k = 2$ group assignments in the discovery and internal assessment arms, respectively. PC1 and PC2 explain 31.7% and 5.6% of the variance. (c,d) Kaplan–Meier curves and numbers at risk for the PFS-like exploratory endpoint in the same two arms. Shading shows 95% confidence intervals. Survival behavior was used when choosing this dataset partition. The log-rank tests are therefore post-selection descriptive results and do not provide independent validation.

survival did not differ (log-rank $p = 0.3130$). Univariable high-versus-low (hazard ratio 1.10, 95% confidence interval 0.91–1.32; $p = 0.3280$) and continuous per-standard-deviation models (hazard ratio 0.98, 95% confidence interval 0.90–1.08; $p = 0.7182$) were nonsignificant. Covariate-complete multivariable models included only 289 samples and were also nonsignificant.

#### Focused EMT and regulator-removal sensitivity analyses

Descriptive single-cell review motivated a post hoc seven-gene EMT-related set:

$$\{ZEB1, ZEB2, VIM, MMP2, COL1A1, COL3A1, TAGLN\}.$$

This set was treated as a familiar EMT and stromal-remodeling program, not as a novel *VIM–ZEB1–ZEB2* pathway.

To test whether the survival-associated structure depended only on the two transcriptional regulators, *ZEB1* and *ZEB2* were removed. The remaining five-gene support set was:

$$\{VIM, MMP2, COL1A1, COL3A1, TAGLN\}.$$

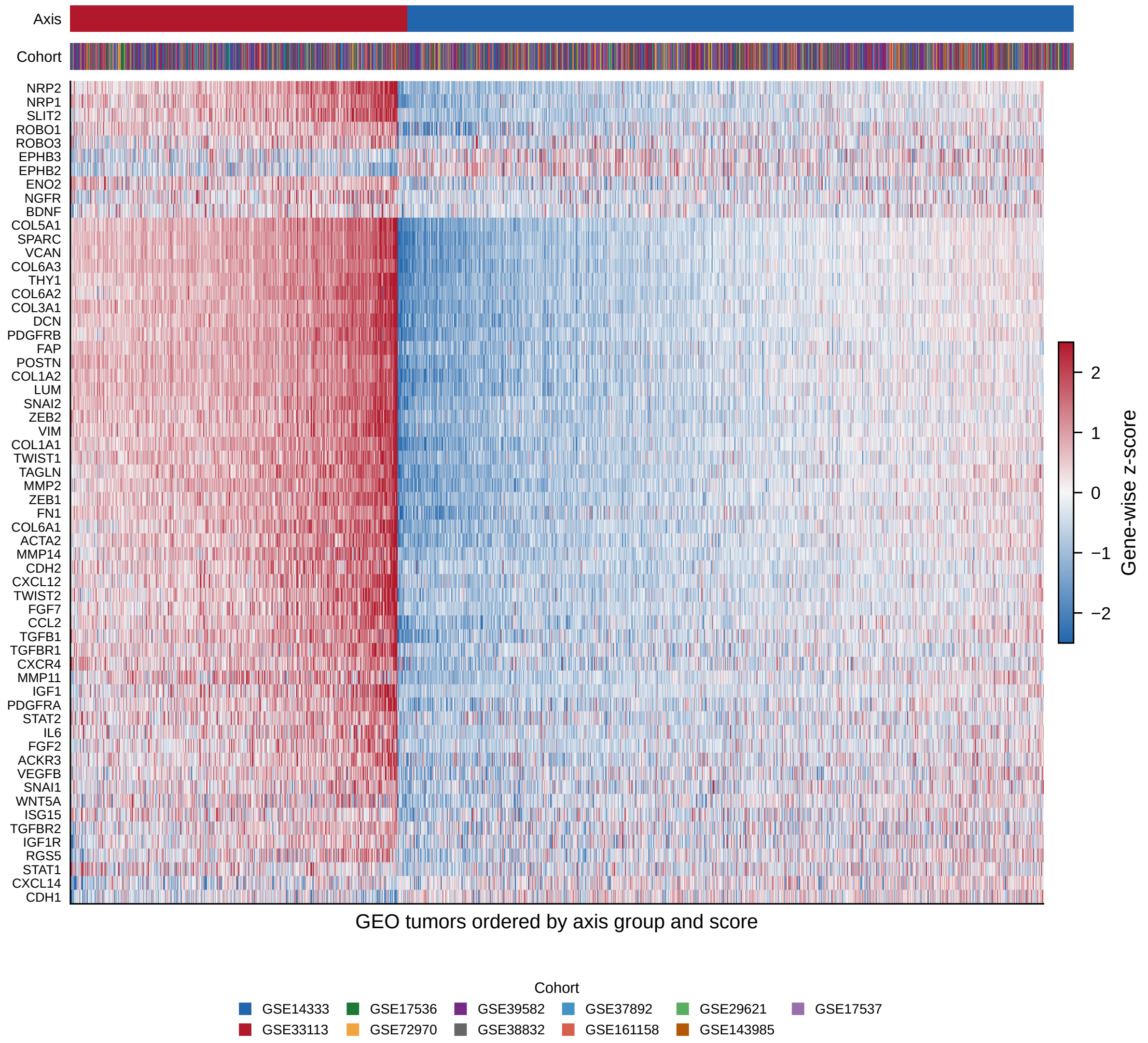


**Figure S3. Selected neural–stromal genes in the GEO modeling set.** Rows are gene-wise standardized expression values; columns are tumors ordered by axis group and score. The display shows 60 axis-associated features. Annotation bars identify axis group and source cohort.

Using the fixed six discovery cohorts and the same preprocessing and clustering procedure, $k = 2$ produced 183 high-risk and 289 low-risk samples (log-rank $p = 6.33 \times 10^{-6}$; source-reported high-to-low incidence ratio 2.60). For $k = 3$, high-, intermediate-, and low-risk groups contained 110, 220, and 142 samples (log-rank $p = 1.38 \times 10^{-4}$; source-reported high-to-low incidence ratio 2.64). Because the gene set was refined after reviewing the data, these results are exploratory.

#### Detailed single-cell and Scissor summaries

The processed GSE178341 object contained 370,115 retained cells and 43,113 genes. All genes in the 84-gene panel were found; 15 low-detection genes were retained. The 84-gene localization score was the unsigned mean of per-gene standardized log-normalized expression. CAFs/fibroblasts, other stromal cells, endothelial cells, and macrophage/myeloid cells had the four highest reported median scores.

The official Scissor Cox pilot sampled 10,000 cells with cell-type balancing. It identified 214 Scissor-positive, 355 Scissor-negative, and 9,431 neutral cells. Table S4 reports positive fractions by broad compartment. These fractions describe the balanced pilot and are not estimates of cell-type prevalence in the original atlas.

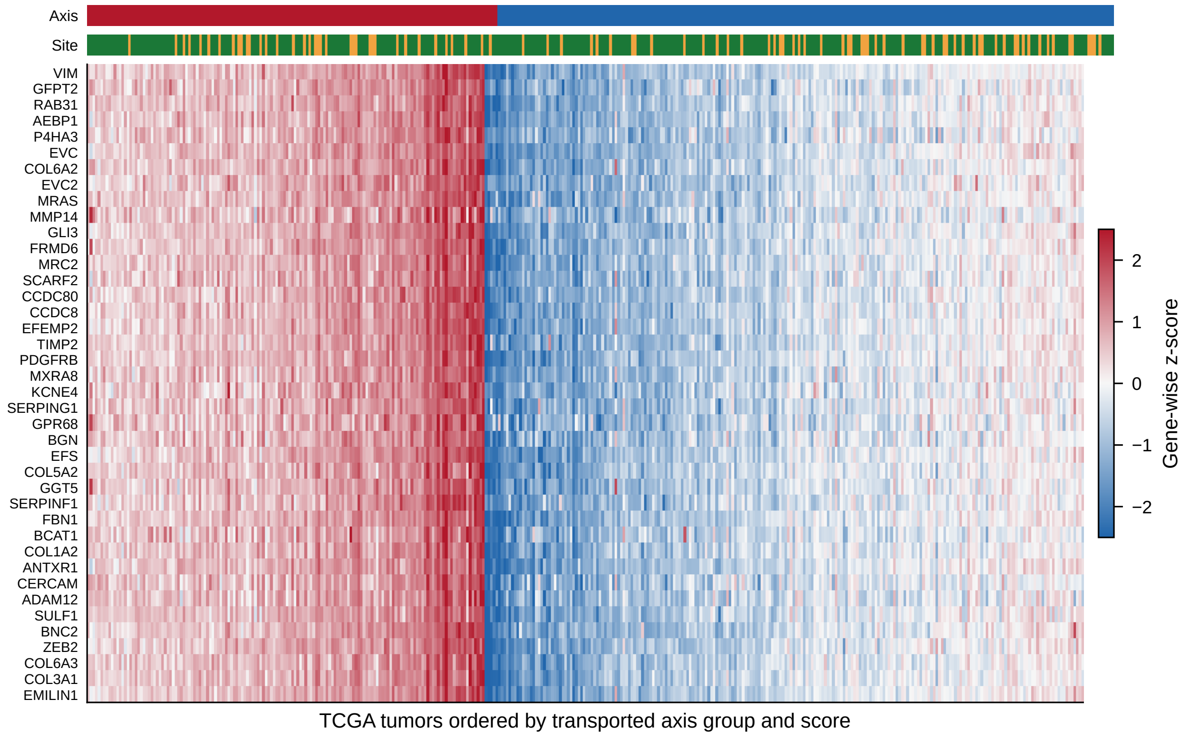


**Figure S4. Top differential genes in TCGA COAD/READ by transported axis group.** Rows are gene-wise standardized expression values across 376 primary tumors ordered by transported group and score. Annotation bars identify axis group and primary cancer site (COAD or READ).

**Table S4. Official Scissor-positive fractions in the balanced 10,000-cell pilot.**

| Broad compartment | Scissor-positive cells (%) |
|---|---|
| Stromal | 7.14 |
| Myeloid | 4.55 |
| Mast | 2.03 |
| Epithelial | 0.56 |
| T/NK/ILC | 0.28 |
| Plasma | 0.21 |
| B cell | 0.21 |

The all-cell Scissor-style approximation used 1,719 bulk samples with 383 events and classified 30,179 cells as positive, 136,015 as negative, and 203,921 as neutral. It omitted the official dense cell–cell network and must not be presented as a full Scissor run.

#### Bulk-cohort comparator and sensitivity results

The complete compact tables are supplied in `supplementary_data/phase3/` through `phase5/`. Their principal roles and results are summarized in Table S5.

#### Independent E-MTAB-12862 external assessment

The E-MTAB-12862 analysis was specified before the survival table was constructed. The official gene-symbol TPM matrix contained 1,063 tumor profiles. Scale inspection without outcome data showed raw TPM-scale expression, which was transformed as $\log_2(\text{TPM} + 1)$. All 87 frozen genes were available, and score construction retained the all-positive, equal-weight definition. Public metadata did not show direct accession overlap with the GEO or TCGA datasets, although patient-level independence cannot be proved without shared cross-study identifiers.

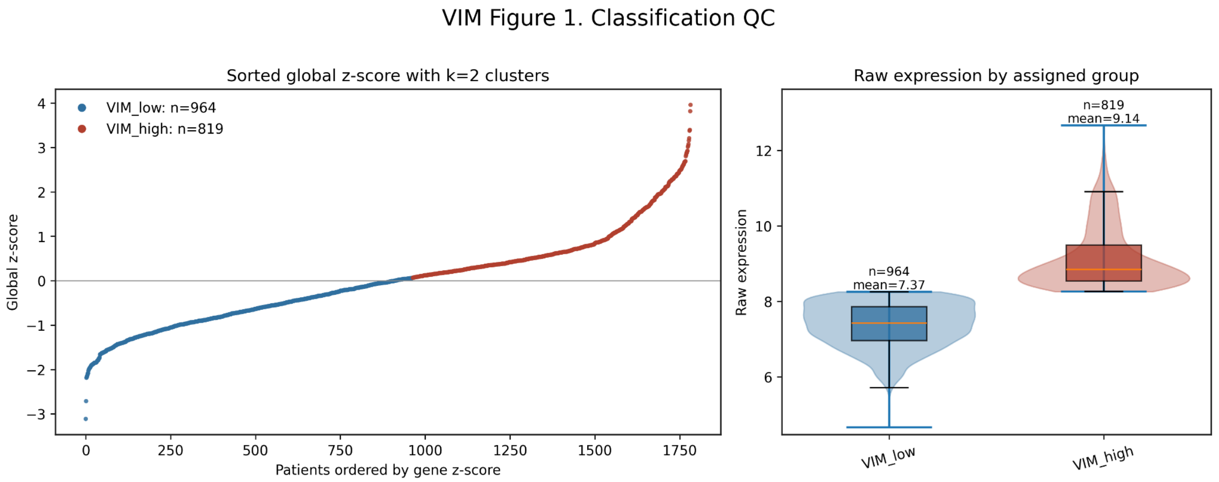


**(a)** Classification quality control and source-scale expression.

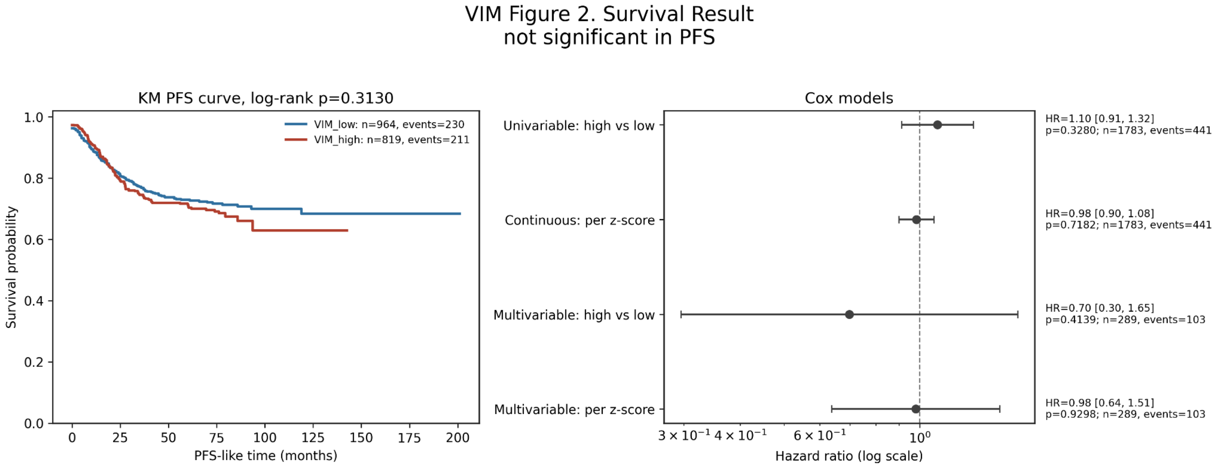


**(b)** Kaplan–Meier and Cox-model results.

**Figure S5. Single-gene *VIM* control.** Expression-based grouping captured the intended *VIM* contrast but did not reproduce the multigene axis survival association.

The source clinical supplement provided RFS and OS. RFS among stage I–III patients was the locked primary endpoint, and OS was secondary. Table S6 reports the continuous-score results. The proportional-hazards test for the axis term gave $p = 0.970$ in the primary RFS model and $p = 0.613$ in the OS model. Stage III did not meet the proportional-hazards assumption in the stage-adjusted RFS model, so that model is interpreted as a sensitivity analysis.

The locked RFS and OS estimates did not reproduce the effect sizes observed in GEO and TCGA. These analyses provide the independent E-MTAB-12862 results for the frozen score.

#### Exploratory model development in E-MTAB-12862

After completing the locked analysis, we used the same 948 stage I–III patients to examine whether outcome-derived gene weights could improve discrimination within E-MTAB-12862. A LASSO-penalized Cox model included standardized expression values for all 87 genes. Five-fold cross-validation across the full RFS analysis set was used to select the penalty. The model was then fitted to all 948 patients and retained 34 genes with nonzero coefficients.

The full-cohort LASSO score was associated with RFS and had an apparent concordance index of 0.658 (Table S7). The largest positive coefficients were assigned to *NRP1*, *CXCL14*, *PDGFRA*, and *ENO2*; the largest negative coefficients were assigned to *THY1*, *SEMA6D*, *ROBO1*, and *CXCR4*. For an internal assessment, coefficients were refitted in each training fold with the previously selected penalty and evaluated in the corresponding held-out fold. The mean held-out concordance index was 0.584. This procedure was not nested because penalty selection used the full E-MTAB-12862

**Table S5. Primary outcome, comparator, and sensitivity analyses.**

| Analysis | Result and inferential role |
|---|---|
| GEO random-effects meta-analysis | 11 cohorts, 1,783 tumors, and 441 events; pooled HR 1.33 per score SD (95% CI 1.19–1.49), with leave-one-cohort-out estimates of 1.31–1.38. |
| Endpoint-family sensitivity | Seven DFS cohorts gave HR 1.39 (1.19–1.62); excluding the singleton RFS, MFS, or metastasis/recurrence-free cohort gave HRs 1.38, 1.33, and 1.32 without direction reversal. |
| Locked TCGA transport | 376 tumors and 102 PFI events; continuous HR 1.25 (1.03–1.52) and locked high-versus-low HR 1.73 (1.17–2.57). |
| CMS4/stroma comparators | Strong score correlations with CMS4-like, CAF, and stroma measures place the axis within the broader mesenchymal phenotype; nested and residualized models did not detect additional prognostic information. |
| Cross-fitted residual axis | On the common original-axis scale, GEO HR 1.12 (0.77–1.64), prediction interval 0.76–1.65; TCGA HR 1.31 (0.63–2.74). Linear and ridge nuisance models agreed. |
| Stage-adjusted sensitivity | Seven cohorts, 943 patients, and 225 events; original-axis HR 1.26 (1.06–1.50) and cross-fitted residual-axis HR 0.94 (0.62–1.41). |
| Module and null controls | Stromal-only HR 1.34; full score minus stromal genes HR 1.10; no leave-one-gene-out reversal; 3 of 1,000 matched random sets were at least as extreme. |

**Table S6. Locked E-MTAB-12862 survival and composition results.**

| Analysis | Result |
|---|---|
| Primary RFS | 948 patients, 436 events; HR 1.05 per score SD (95% CI 0.95–1.16; $p = 0.346$) |
| Stage-adjusted RFS | 948 patients, 436 events; axis HR 1.03 (0.93–1.14; $p = 0.525$) |
| Secondary OS | 1,062 patients, 502 deaths; HR 1.06 (0.97–1.16; $p = 0.193$) |
| Stromal-remodeling module | RFS HR 1.05 (0.95–1.16) |
| Neural-guidance module | RFS HR 0.98 (0.89–1.08) |
| GEO-trained residual axis | RFS HR 1.001 (0.998–1.004) on the non-restandardized transported scale |
| Composition correlations | CMS4 binary label $\rho = 0.699$; CAF proxy $\rho = 0.953$; stroma proxy $\rho = 0.954$; purity proxy $\rho = -0.909$ |
| *VIM* control | RFS HR 1.06 (0.96–1.18; $p = 0.239$) |

outcome dataset.

These estimates do not provide external validation; the fixed 34-gene score requires assessment in another cohort.

**Table S7. Frozen-score external assessment and exploratory model development in E-MTAB-12862.**

| Analysis | Result and interpretation |
|---|---|
| Frozen 87-gene score | 948 patients and 436 RFS events; HR 1.05 per score SD (95% CI 0.95–1.16); apparent concordance index 0.513. This was the locked independent analysis. |
| Full-cohort LASSO fit | 34 genes; HR 1.80 per score SD (1.64–1.98); apparent concordance index 0.658. The same RFS data were used to select and estimate the model. |
| Fold-specific assessment | Mean concordance index 0.584 (SD 0.023) across five held-out folds. The penalty was selected once from the full cohort, so this was not nested cross-validation. |
| Highest versus lowest quartile | 237 patients per group, with 172 and 66 events, respectively; HR 4.07 (3.06–5.42). This is a descriptive comparison within the model-development cohort. |

#### Simulation benchmark of composition residualization

The corrected benchmark used the same common effect scale as the primary cohort analysis: log hazard ratio per one within-cohort standard deviation of the original axis, without method-specific residual re-scaling. Thirteen scenarios, six methods, and 500 final repetitions produced 39,000 pooled fits. All pooled fits completed. Of 429,000 cohort-level fits, 472 failed and remained in the reported denominators. Table S8 summarizes prespecified cross-fitted linear-

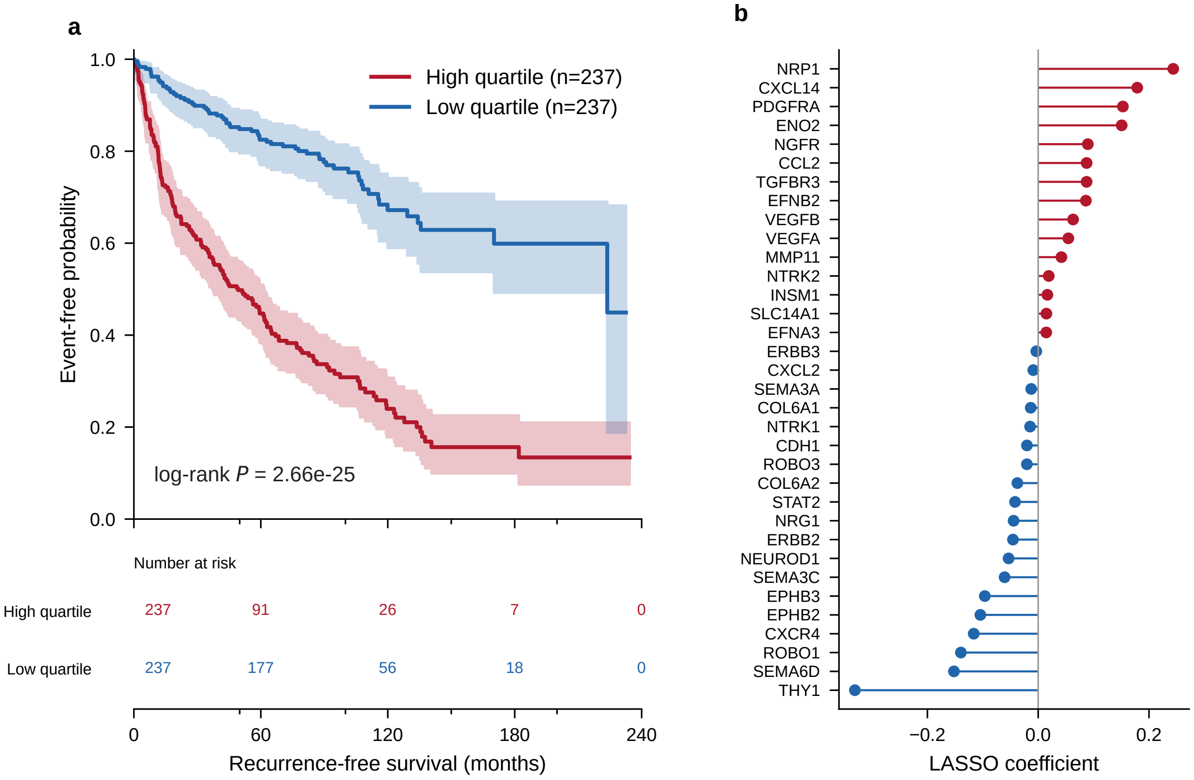


**Figure S6. Exploratory E-MTAB-12862 LASSO model.** (a) Kaplan–Meier curves compare the highest and lowest score quartiles; the middle 50% of patients are not included. Shading shows 95% confidence intervals. (b) Nonzero coefficients for the 34 genes retained by the model; positive coefficients indicate higher fitted hazard. The score, penalty, and coefficients were estimated in the same 948 patients used for panel a. This same-cohort result does not provide external validation.

residual results; complete results for all six methods are in `supplementary_data/work_package_f/`.

**Table S8. Selected cross-fitted linear-residual simulation results.** Coverage is against the clean biological target except where an observed-endpoint value is also shown. FPR, false-positive rate.

| Scenario | FPR | Adverse power | Clean coverage | Observed coverage |
|---|---|---|---|---|
| Ideal oracle-composition null | 0.038 | – | 0.962 | 0.962 |
| Cohort-shift null | 0.050 | – | 0.948 | 0.948 |
| Noisy-comparator null | 0.102 | – | 0.910 | 0.910 |
| Nonlinear-composition null | 0.248 | – | 0.776 | 0.776 |
| 10% endpoint-noise null | 0.042 | – | 0.960 | 0.960 |
| 10% endpoint-noise small signal | – | 0.446 | 0.926 | 0.948 |

For the 10% endpoint-noise small-signal scenario, the clean target was 0.161 and the observed-endpoint target was 0.131. This difference is not a software error: label noise changes the association estimable from the observed endpoint. The ideal-null calibration supports the implementation, whereas the noisy and nonlinear scenarios show that cross-fitting alone cannot remove residual confounding from poor nuisance measurements or model misspecification.

#### Patient-level single-cell analysis

The corrected patient–tissue–cell-type pseudobulk analysis retained the patient as the inferential unit and analyzed GSE178341 and GSE132465 separately. Five official GEO files were restored from frozen URLs and passed byte-count and MD5 checks. All 370,115 GSE178341 and 63,689 GSE132465 matrix barcodes matched official metadata. All 187 GSE132465 mast cells were assigned to `mast_cell`, with none assigned to T/NK/ILC. At the primary

50-cell threshold, epithelial, myeloid, and broad other-stromal compartments had concordant tumor-higher 84-gene signed state scores where estimable. The GSE178341 fibroblast/CAF state was also tumor-higher but was based on five matched pairs (median difference 0.249, $p = 0.0625$); fibroblast abundance was higher in normal tissue ($p = 0.000398$). Corrected mapping, tumor–normal, abundance, threshold, and leave-one-patient-out tables are in `supplementary_data/work_package_c/`. These data support localization and tumor–normal state differences, not prognosis.

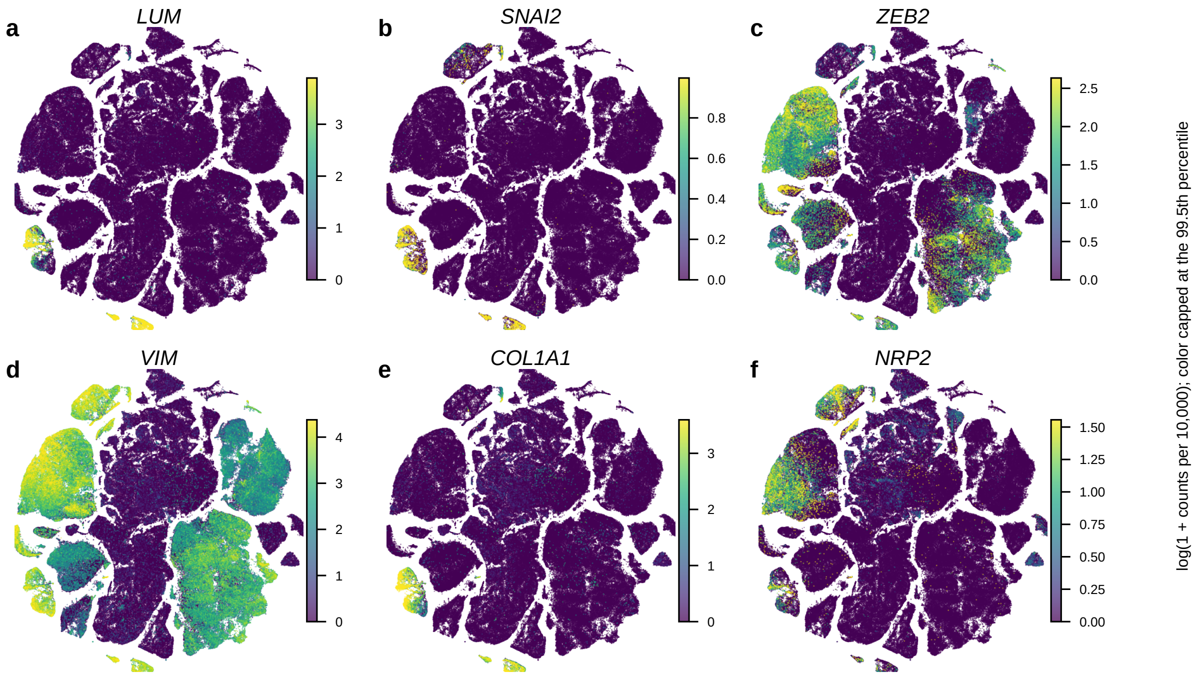


**Figure S7. Representative axis-related genes on the GSE178341 global t-SNE.** (a–f) Cell-level log(1 + counts per 10,000) expression for *LUM*, *SNAI2*, *ZEB2*, *VIM*, *COL1A1*, and *NRP2*. Each panel uses its own expression scale capped at the gene-specific 99.5th percentile. These maps show cellular localization and are not patient-level tests.

#### Spatial projection in GSE267401

Four Visium CytAssist sections from two patients retained 16,171 spots after section-specific quality control. Every primary score feature was detected in every section. Table S9 summarizes the frozen-score projection; complete correlation, Moran's $I$, score-version, and QC-sensitivity tables are in `supplementary_data/phase7/`.

**Table S9. Section-level spatial projection summary.** CAF and stroma proxies are not gene-disjoint from the frozen score.

| Section | Spots | CAF $\rho$ | Stroma $\rho$ | Moran's $I$ | Neural–stroma $\rho$ |
|---|---|---|---|---|---|
| CTC21P | 4,434 | 0.852 | 0.855 | 0.655 | 0.488 |
| CTC21M | 3,687 | 0.895 | 0.915 | 0.730 | 0.348 |
| CTC17P | 3,534 | 0.806 | 0.819 | 0.610 | 0.316 |
| CTC17M | 4,516 | 0.651 | 0.681 | 0.242 | 0.077 |

All 11 original CAF-proxy and 14 original stroma-proxy genes were members of the frozen score, so those correlations are descriptive and partly expected by construction. A gene-disjoint sensitivity removed every candidate marker found in either the 87- or 84-gene list and retained 16 CAF and 15 stromal markers after detection filtering. The frozen score remained positively correlated with both disjoint proxies in all four sections: CAF $\rho = 0.227$, 0.635, 0.814, and 0.765; stroma $\rho = 0.373$, 0.660, 0.808, and 0.722 for CTC17M, CTC17P, CTC21M, and CTC21P, respectively. The attempted block-permutation procedure was not accepted, and all corresponding $p$-value fields are retained as NA. The four sections still represent only two patients, and spot-level replication cannot substitute for patient-level replication.

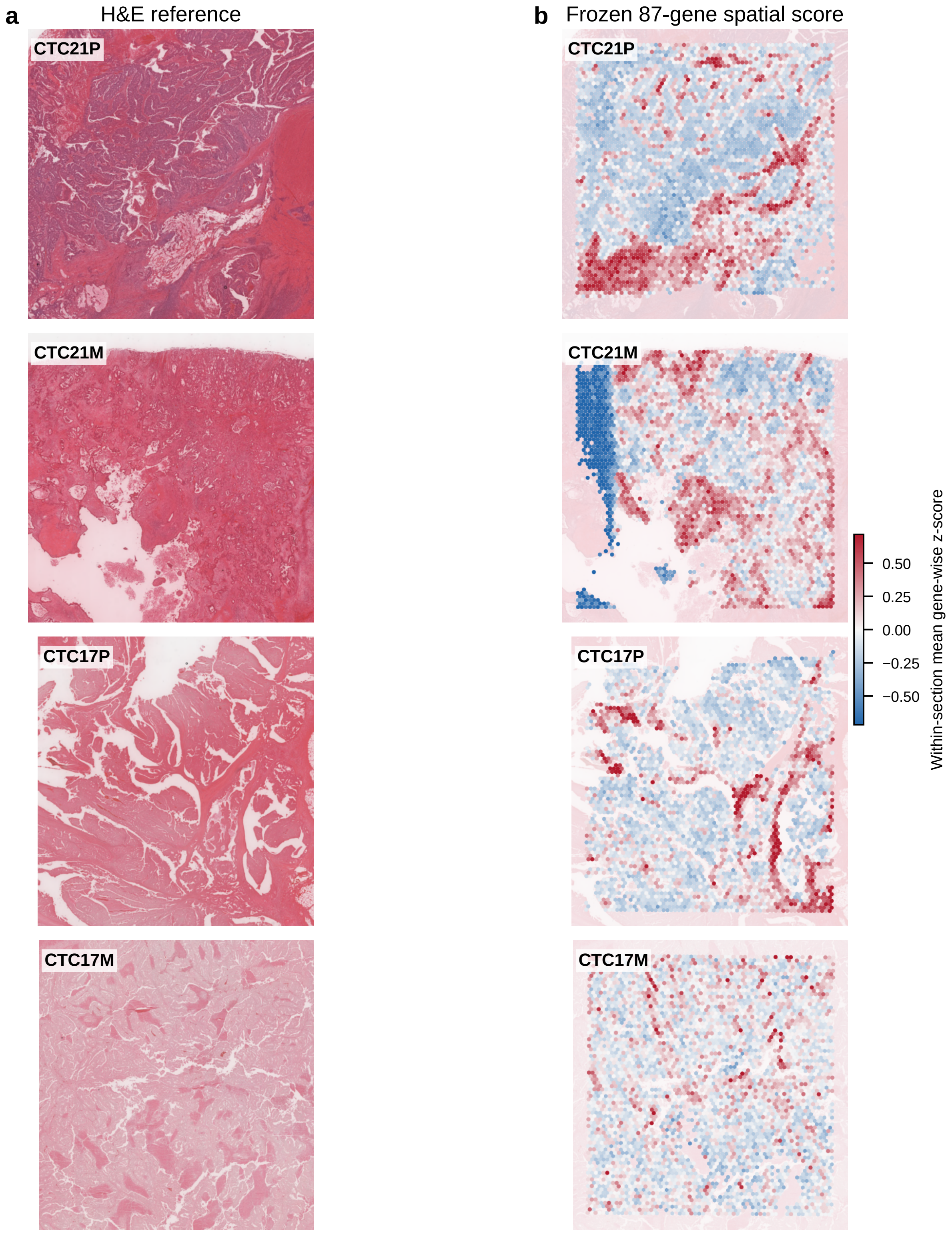


**Figure S8. Histology and frozen 87-gene score in four GSE267401 sections.** (a) H&E reference images. (b) The frozen 87-gene score overlaid on the corresponding tissue image. Scores are means of within-section gene-wise z-scores and use a common color scale across sections. The maps are descriptive: the four sections come from two patients and do not provide a patient-level progression test.

#### Ligand–receptor negative screen

The prespecified expression screen identified 12 database pairs with minimal other-stromal ligand and epithelial receptor expression support in both single-cell datasets: *BDNF–NTRK2*, *CDH1–CDH1*, *CDH2–CDH2*, *CXCL12–ACKR3*, *CXCL12–CXCR4*, *EFNB2–EPHB2*, *EFNB2–EPHB3*, *EFNB2–EPHB4*, *IGF1–IGF1R*, *NCAM1–NCAM1*, *NGF–NGFR*, and *NGF–NTRK1*. None met the combined patient-expression, spatial, bulk-outcome, and prior-evidence criteria. The initially screened *SEMA3C–NRP1_NRP2* row was excluded because the multi-subunit receptor was handled inconsistently. No primary interaction was selected. Machine-readable replication, detection, evidence, and sender–receiver tables are in `supplementary_data/phase8/`. This is a pseudobulk expression screen, not formal communication-probability inference.

#### Estimand-and-evidence map

**Table S10. Estimand and evidence map for the principal analyses.**

| Analysis | Population, estimand, and method | Permitted interpretation |
|---|---|---|
| Primary GEO meta-analysis | Patient-level analysis of 1,783 tumors and 441 events from 11 GEO cohorts. Separate Cox models estimated the hazard ratio per within-cohort standard-deviation increase in the frozen 87-gene score using each study's source progression-related endpoint. Log hazard ratios were pooled by REML with Knapp–Hartung uncertainty. | Association between the frozen score and progression-related survival across the included GEO studies |
| Locked TCGA transport | Patient-level analysis of 376 TCGA COAD/READ primary tumors with 102 PFI events. The continuous score was standardized within TCGA. The frozen GEO scaler and $k$-means centroids were used only for high- and low-axis assignment. Both exposures were calculated before PFI outcomes were joined. | External support for the direction and transportability of the association, not prospective clinical validation |
| Cross-fitted residual analysis | Patient-level analysis of 11 GEO cohorts. A model of the axis based on CMS4-like affinity, CAF, stroma, and purity was trained in ten cohorts and applied to the held-out cohort. Outcomes were not used to construct the residual, which remained on the original-axis scale. Cohort Cox estimates were pooled by REML with Knapp–Hartung uncertainty. | Detectable association beyond *measured composition under the stated nuisance model*; this does not establish removal of unmeasured or poorly measured composition |
| Stage-adjusted sensitivity | Patient-level analysis of seven stage-eligible GEO cohorts, including 943 patients and 225 events. Cohort-specific Cox models included categorical stage with either the original score or the cross-fitted residual; estimates were pooled by REML with Knapp–Hartung uncertainty. | Sensitivity among the patients and cohorts with usable stage, not evidence of clinical independence across all included studies |
| Single-cell pseudobulk analysis | Patient–tissue–broad-cell-type pseudobulk profiles from GSE178341 and GSE132465, representing 62 and 23 patients. Paired patient-level tests estimated tumor–normal differences in the 84-gene signed state score where matched samples were available; unpaired tests were labeled exploratory. | Cellular localization and tumor-associated state differences, not single-cell prognostic validation |
| Spatial sensitivity analysis | Section-level analysis of four sections from two patients, with 16,171 spots retained after quality control. Within-section rank correlations compared the frozen score with gene-disjoint CAF and stromal proxies. Patient summaries were descriptive. | Descriptive spatial concordance in two patients, not population-level inference, independent validation, or evidence of progression association |
| Simulation benchmark | Synthetic repetitions preserved the sizes and event structure of the 11 GEO cohorts. Thirteen frozen scenarios compared six analysis methods in 500 repetitions per scenario–method combination, producing 39,000 pooled meta-analysis fits. Performance was measured against known simulation targets. | Operating characteristics under the specified simulated assumptions, not proof that the assumptions or unbiasedness hold in the real patient data |

#### Machine-readable supplementary data

The `supplementary_data/` directory contains the exact gene definitions, cohort and endpoint maps, source-code provenance, and compact result tables for the bulk, single-cell, spatial, ligand–receptor, simulation, E-MTAB-12862 external-assessment, and exploratory model-development analyses. Its `README.md` maps each file to the corresponding manuscript result. The single-cell analysis was rerun from checksum-verified GEO files. The spatial directory reports section correlations and patient-level descriptive summaries; fields from the withdrawn block-permutation analysis remain NA and are not used in the manuscript. Raw public data and large intermediate matrices are not duplicated in the repository, and the documented restoration command retrieves and verifies the required single-cell inputs.

#### Computational workflow summary

**Table S11. Major computational components and their inferential roles.**

| Component | Methods | Role and interpretation |
|---|---|---|
| GEO preprocessing | Probe-to-gene mapping, mean probe collapse, within-dataset standardization, clinical matching | Create comparable GPL570 gene-level matrices, retain each source endpoint, and standardize time and event fields for the exploratory PFS-like display. |
| Axis grouping | Evidence-gene PCA; $k$-means with $k = 2$, plus $k = 3$ sensitivity analyses | Define expression groups without directly fitting individual survival times during clustering; dataset partition selection was nevertheless outcome informed. |
| Survival | Kaplan–Meier curves, log-rank tests, Cox proportional-hazards models | Describe the selected GEO split and estimate standardized-score associations separately within each source endpoint before meta-analysis. |
| Endpoint sensitivity | DFS-only and leave-one-endpoint-family-out REML meta-analyses | Test whether the pooled direction depends on one non-DFS endpoint family without treating singleton families as meta-analyses. |
| Differential expression | `limma` empirical-Bayes models and Benjamini–Hochberg FDR | Characterize high- versus low-axis transcriptional differences. |
| Pathways | Preranked gene-set enrichment and hypergeometric over-representation tests | Interpret Hallmark, GO, KEGG, and custom module signals. |
| Microenvironment | EPIC deconvolution, marker z-scores, Wilcoxon tests, rank-biserial effects | Estimate cellular composition and exploratory cell-state programs in bulk tumors. |
| Locked TCGA transport | Transported GEO scoring/orientation to TCGA; PFI survival and expression analyses | Assess outcome direction and molecular concordance without TCGA outcome tuning. |
| Independent E-MTAB external assessment | Locked 87-gene scoring in 1,063 tumors; primary stage I–III RFS model; secondary OS, stage, module, composition, residual-axis, and *VIM* analyses | Test the frozen score in a separate population-based RNA-sequencing cohort without outcome-informed changes to membership, direction, scale, or cutoff. |
| Exploratory E-MTAB model development | LASSO-penalized Cox regression using the 87 genes; full-cohort fit, descriptive quartile comparison, and fixed-penalty fold assessment | Examine outcome-derived weighting within E-MTAB-12862. These analyses use the same RFS data for model development and do not constitute external validation. |
| Single-cell projection | Counts-per-10,000 normalization, log transformation, per-gene standardization, mean and weighted scores | Locate the bulk-derived axis across GSE178341 cell annotations. |
| Bulk–single-cell association | Official Scissor Cox pilot; all-cell Cox score-statistic approximation | Identify phenotype-associated cells; only the pilot uses the official Scissor network method. |
| CMS4/stroma relationship | Comparator correlations, nested Cox models, common-scale study-level cross-fitted residual scores, and categorical-stage sensitivity | Test whether the axis contributes information beyond established mesenchymal and stromal signals without using held-out outcomes in residual construction, and assess the original and residual scores in stage-eligible cohorts. |
| Simulation benchmark | Thirteen frozen scenarios, six methods, 500 repetitions, explicit oracle targets, and retained fit failures | Verify ideal-null calibration and identify failure modes caused by noisy comparators, nonlinear nuisance structure, and endpoint-label error. |
| Module controls | Module ablation, leave-one-gene-out analysis, and expression-matched random gene sets | Identify the dominant component and benchmark specificity against comparable expression sets. |
| Patient-level single cell | Checksum-verified public-input restoration, exact barcode matching, corrected mast-cell mapping, patient–tissue–cell-type pseudobulk scoring, paired contrasts, threshold sensitivity, and abundance models | Reproduce the analysis from official GEO files and separate within-compartment state from cell abundance without treating cells as replicates. |
| Spatial projection | Section-wise frozen scoring, gene-disjoint CAF/stromal sensitivity, rank correlations, Moran's $I$, neighborhoods, and QC sensitivity | Assess descriptive spatial organization while disclosing that only two independent patients are available. |
| Ligand–receptor screen | Prespecified sender–receiver expression replication and multi-source evidence filter | Retain a negative result; co-expression alone is insufficient to infer communication. |
